# The transcription factor THAP11 is a general regulator of early embryonic polarization

**DOI:** 10.64898/2026.09.21.753235

**Authors:** Qianying Yang, Shan Jiang, Boyan Wang, Yi Zhang

## Abstract

During mouse embryogenesis, three sequential waves of cell polarization occur in the 8-cell-morula trophectoderm (TE), E3.5-E4.5 primitive endoderm (PrE), and E4.5-E5.5 epiblast (Epi), driving cell fate determination and morphogenic events that lay the foundation for embryonic development. However, it is not clear whether a general regulator that functions in all three waves of polarization exist. Here, we demonstrate that transcriptional factor THAP11 is such a general regulator. Using a novel mouse model combining genetic knockout and an inducible degron system, we demonstrate that stage-specific acute THAP11 depletion causes the failure of the three waves cell polarization, resulting in defective blastocyst formation, PrE maturation, and Epi lumenogenesis, respectively. Mechanistically, THAP11 does not directly regulate pluripotency genes but drive polarization through both stage-specific and conserved chromatin-bindings that commonly activate polarity programs, including Golgi vesicle transport. THAP11 activates polarity genes by promoting chromatin accessibility at their promoters. THAP11, together with stage-specific regulators, sequentially paces the timing of cell polarization. Collectively, our study establishes THAP11 as a general regulator of embryonic morphogenesis.

## Main

Cell polarity establishment is a major driver of embryonic morphogenesis ^1,2^. It occurs at defined developmental stages in a spatial-temporally regulated manner to promote lineage segregation and increase embryonic structural and functional complexity. During early embryonic development, mouse embryos undergo the first cell polarization from 8-cell to morula stages, driving the first cell fate decision to generate the apolar inner cell mass (ICM) and the polarized extraembryonic trophectoderm (TE) ^2–4^. The ICM cells undergo the second wave of polarization from embryonic day 3.5 (E3.5) to E4.5, resulting in lineage segregation of the apolar naïve epiblast (Epi) and the polarized primitive endoderm (PrE) ^2,5,6^. During implantation, naïve Epi cells initiate polarization and lumenogenesis as they transition to formative Epi from E4.5 to E5.5, thereby acquiring competence to differentiate into all somatic and germline lineages ^7–9^. Despite the essential roles of these sequential polarization events in early embryonic development, how they are spatial-temporally regulated remains elusive.

Developmental stage-specific studies of cell polarity establishment have identified transcriptional activators TFAP2C and TEAD4 in TE polarization ^10^, and transcriptional repressors OCT4 and SOX2 in Epi polarization ^11,12^. However, these factors do not explain how cell polarization is sequentially regulated during early embryogenesis. Cell polarization is a highly conserved and organized process involving Golgi network organization, GTPase signal transduction, and actin cytoskeleton remodeling ^13,14^, suggesting the existence of a common regulator that coordinates these regulations. We hypothesis that such general transcriptional regulator should activate key polarity programs, acting in concert with stage-specific regulators to coordinate these sequential polarization events. Consistent with this hypothesis, core polarity components, including protein kinase C (PKC) and partitioning-defective (PAR) molecules ^4–6,8,15^ are required across these stages.

In this study, we identify and demonstrate that the transcription factor THAP11 is a general regulator for early embryonic polarization. Using novel mouse models that combine genetic knockout and targeted protein degradation tag (dTAG) system ^16,17^, together with RNA-seq, ATAC-seq, and low-input CUT&RUN analyses, we provide multiple pieces of evidence supporting THAP11’s role in regulating all three cell polarization stages. Phenotypically, stage-specific depletion of THAP11 disrupts all three waves of cell polarization during early embryogenesis, resulting in failure of blastocyst formation, PrE maturation, and Epi lumenogenesis, respectively. Mechanistically, THAP11 does not directly regulate pluripotency genes, but preferentially binds to promoter regions of polarity genes across all three waves of polarization. Notably, both stage-specific and conserved THAP11 chromatin-binding play an essential role in regulating expression of polarity genes, including those involved in actin filament organization, GTPase signal transduction, and Golgi vesicle transport, by promoting chromatin accessibility at their promoters. Together, our findings not only establish THAP11 as a general regulator of all three waves cell polarization in early embryogenesis, but also advance our understanding of how these sequential polarization events are coordinately regulated by a single transcription factor.

## Results

### Identification of THAP11 as a potential regulator for all three waves of polarization

To identify potential regulators for all three waves of cell polarization during early embryonic development, we first generated a list of 2,285 cell polarity-related genes from public datasets (**Supplementary Table 1**). Of this gene list, 1,133 core cell polarity-related genes are expressed in all three polarized cell types: morula, E4.5 PrE, and E5.5 Epi (**Supplementary information, Fig. S1a**). Motif enrichment analysis of open chromatin regions at promoters of these core cell polarity genes, combined with Homer and ChIP-Atlas analyses revealed 7 candidate regulators (**Fig. 1a and Supplementary information, Fig. S1a**). In contrast to known stage-specific regulators whose motifs and expression are enriched at particular stages, such as GATA4 in E4.5 PrE and SOX2 in E5.5 Epi, these 7 candidates were expressed and enriched across all three polarization stages (**Fig. 1a and Supplementary information, Fig. S1b**). Of the 7 candidates, only 3 (*Gabpa, Ctcf, Thap11*) exhibit embryonic lethal phenotype before E6.5 (**Fig. 1a**). Since previous studies have shown that GABPA 17 and CTCF 18 are dispensable for TE formation and PrE maturation, indicating normal TE and PrE polarization, we therefore focused on THAP11 (also known as RONIN). Although *Thap11* zygotic knockout embryos could develop to blastocyst stage 19, potential functional compensation by maternally deposited THAP11 (**Supplementary information, Fig. S1c**) could be an explanation.

**Fig. 1.**
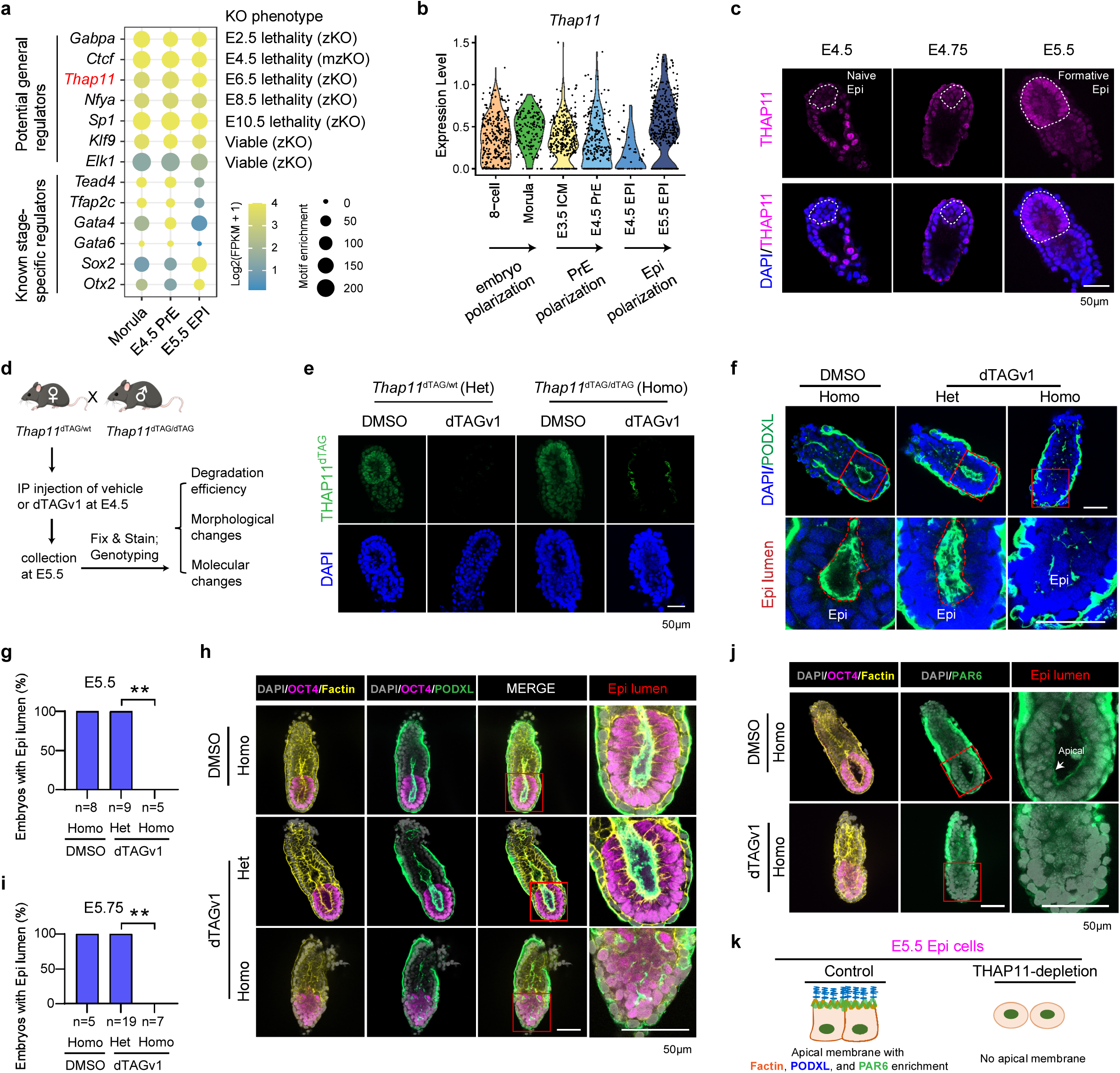
Identifying THAP11 as a potential regulator of cell polarization and its depletion causes epiblast morphogenesis failure. **a**, Left panel: TF motif enrichment at the ATAC–seq peaks of expressed core polarity gene promoters at morula, E4.5 PrE, and E5.5 Epi. Right panel: summary of KO phenotypes. b, Dynamics of *Thap11* expression during the three waves of early embryo cell polarization. c, Immunostaining of THAP11 in embryos during Epi polarization. The white dashed circle marks the epiblast cells. d, Schematic diagram of experimental design. e, Immunostaining of THAP11 in Het and Homo E5.5 embryos under DMSO or dTAGv1 treatment. f, Immunostaining of PODXL in Het and Homo E5.5 embryos under DMSO or dTAGv1 treatment. A high-resolution view of the boxed regions is shown in the bottom panels. The red dashed circle marks the epiblast lumen. g, The ratio of embryos with Epi lumen. Chi-squared test, **p<0.01. h, Immunostaining of OCT4, F-actin, and PODXL in Het and Homo E5.75 embryos under DMSO or dTAGv1 treatment. A high-resolution view of the boxed regions is shown on the right. i, The ratio of embryos with Epi lumen. Chi-squared test, **p<0.01. j, Immunostaining of OCT4, F-actin, and PAR6 in Homo E5.75 embryos under DMSO or dTAGv1 treatment. A high-resolution view of the boxed regions is shown on the right. The white arrow indicates the PAR6 expression on apical membrane of epiblast cells. k, A diagram summarizing the defects of THAP11-depleted E5.5 Epi cells. Scale bar, 50 μm.

Consistent with the notion that THAP11 may serve as a general regulator of these three waves cell polarization, its transcription is increased during 8-cell to morula polarization, was maintained at relatively higher levels in PrE than E4.5 Epi cells, and increased during E4.5 to E5.5 Epi polarization (**Fig. 1b**). Immunostaining further confirmed a increase during E4.5 to E5.5 Epi polarization (**Fig. 1c**). Thus, we first focused our effort on THAP11’s role in Epi polarization.

### dTAG-based THAP11 degradation impairs epiblast morphogenesis

To directly address THAP11’s function during E4.5 to E5.5 Epi morphogenesis, we utilized a rapid protein degradation dTAG system as its successful application in mouse embryos ^16,17^ allows the stage-specific role of THAP11 in cell polarization to be addressed. To this end, we designed a C-terminal THAP11 - FKBP (FK506 binding proteins) - HA (hemagglutinin) fusion construct (referred to as THAP11^dTAG^) to enable conditional degradation by dTAG compounds (*eg.*, dTAG13 and dTAGv1) (**Supplementary information, Fig. S1d**). After confirming the knock-in of the fusion protein and the efficiency of dTAG13-induced degradation in mouse ES cells (mESCs) (**Supplementary information, Fig. S1e**), we tested THAP11’s role in Epi polarization using an *in vitro* 3D culture system ^8^ (**Supplementary information, Fig. S1f**). Immunostaining showed that control spheroids formed a centralized apical membrane with organized F-actin, accompanied by expression of the epithelial polarity marker PAR6 and the lumenogenesis marker podocalyxin (PODXL). In contrast, THAP11-depletion resulted in failure of cell polarization, characterized by disorganized F-actin and loss of PAR6 and PODXL expression (**Supplementary information, Fig. S1g-h**), suggesting an essential role of THAP11 in self-organization, cell polarity establishment, and lumenogenesis of *in vitro* epiblast cells. We then generated the *Thap11*^dTAG^ mouse line using the CRISPR technology ^20^ (**Supplementary information, Fig. S1i**). Notably, these mice are fertile and developmentally normal, with an average litter-size of 9.5 pups based on 12 litters obtained from *Thap11*^dTAG/dTAG^ intercrosses (**Supplementary information, Fig. S1j**), comparable to WT controls (**Supplementary information, Fig. S1k**), indicating that the dTAG knock-in does not affect THAP11 function under physiological conditions. Importantly, dTAGv1 treatment of cultured *Thap11*^dTAG^ E.5 embryos can efficiently deplete THAP11 within 1h (**Supplementary information, Fig. S1l**). Collectively, these results demonstrate that we have successful generated a THAP11^dTAG^ mouse model.

To evaluate the role of THAP11 during E4.5 to E5.5 Epi morphogenesis, we first tested its *in vivo* degradation efficiency (**Fig. 1d**), which showed that THAP11^dTAG^ protein (labeled by FKBP-HA) can be completely depleted in both *Thap11*^dTAG/wt^ (Het) and *Thap11*^dTAG/dTAG^ (Homo) embryos by IP injection of dTAGv1 (**Fig. 1e**). We next examined their morphological changes and observed strong PODXL expression in Epi lumen of E5.5 DMSO-Homo and dTAGv1-Het embryos, whereas PODXL expression and Epi lumen were lost in E5.5 THAP11-depleted (dTAGv1-Homo) embryos (**Fig. 1f-g**), suggesting that the THAP11-depleted embryos failed to undergo Epi lumenogenesis. To exclude the possibility that the observed defects reflect developmental delay, we collected E5.75 embryos. Immunostaining revealed that only THAP11-depleted embryos lost Epi lumen and exhibited disorganized F-actin (**Fig. 1h-i**), indicating a failure of cell polarization. Consistently, apical PAR6 localization was detected in lumen side of Epi cells from control embryos but was absent upon THAP11-depletion (**Fig. 1j and Supplementary information, Fig. S1m**), demonstrating a critical role of THAP11 in Epi polarization. Collectively, the above results indicate that THAP11 is required for polarization and lumenogenesis of Epi cells during peri-implantation development (**Fig. 1k**).

### Acute THAP11 loss does not affect pluripotency transition both *in vivo* and *in vitro*

To understand how THAP11 regulates Epi cell polarization, we performed RNA-seq using E5.5 Epi cells (**Fig. 2a and Supplementary information, Fig. S2a**). The purity of the Epi cells was confirmed by the expression of lineage marker genes (**Supplementary information, Fig. S2b**). To unbiasedly assess the impaired biological processes in THAP11-depleted Epi cells, we performed comparative gene set enrichment analysis (GSEA) of the DMSO, dTAGv1-Het, and dTAGv1-Homo samples. We found that genes involved in cell polarity establishment were significantly downregulated in dTAGv1-Homo relative to both DMSO and dTAGv1-Het samples (**Fig. 2b**), consistent with the observed polarization defects (**Fig. 1f-j**). As previous long-term THAP11 knockout impairs pluripotency in mESCs ^21^ and the exit from naïve pluripotency is required for cell polarization in Epi ^22^, we were wondering whether polarization defects in THAP11-depleted Epi are caused by failure of pluripotency transition. To this end, we systematically analyzed the expression of naïve and formative pluripotent genes upon THAP11 depletion *in vivo* and the results showed no significant changes in the pluripotency marker genes (**Supplementary information, Fig. S2c**), indicating normal progression of naïve-formative transition upon THAP11 depletion. We also performed a similar analysis in 3D cultured ESCs (**Supplementary information, Fig. S2d-e**) and 2D cultured ESCs (**Supplementary information, Fig. S2f-h**). Both results showed no significant changes of naïve and formative marker genes upon THAP11-depletion during naïve-formative transition (**Supplementary information, Fig. S2e, g-h**). Notably, our acute depletion results are different from the previous long-term knockout results by Salewskij et al. ^21^, which showed that long-term *Thap11*-KO in ESCs leads to upregulation of naïve marker genes and downregulation of formative marker genes. Taken together, our results demonstrate that acute THAP11 depletion does not impair naïve-to-formative pluripotency transition both *in vivo* and *in vitro*.

**Fig. 2.**
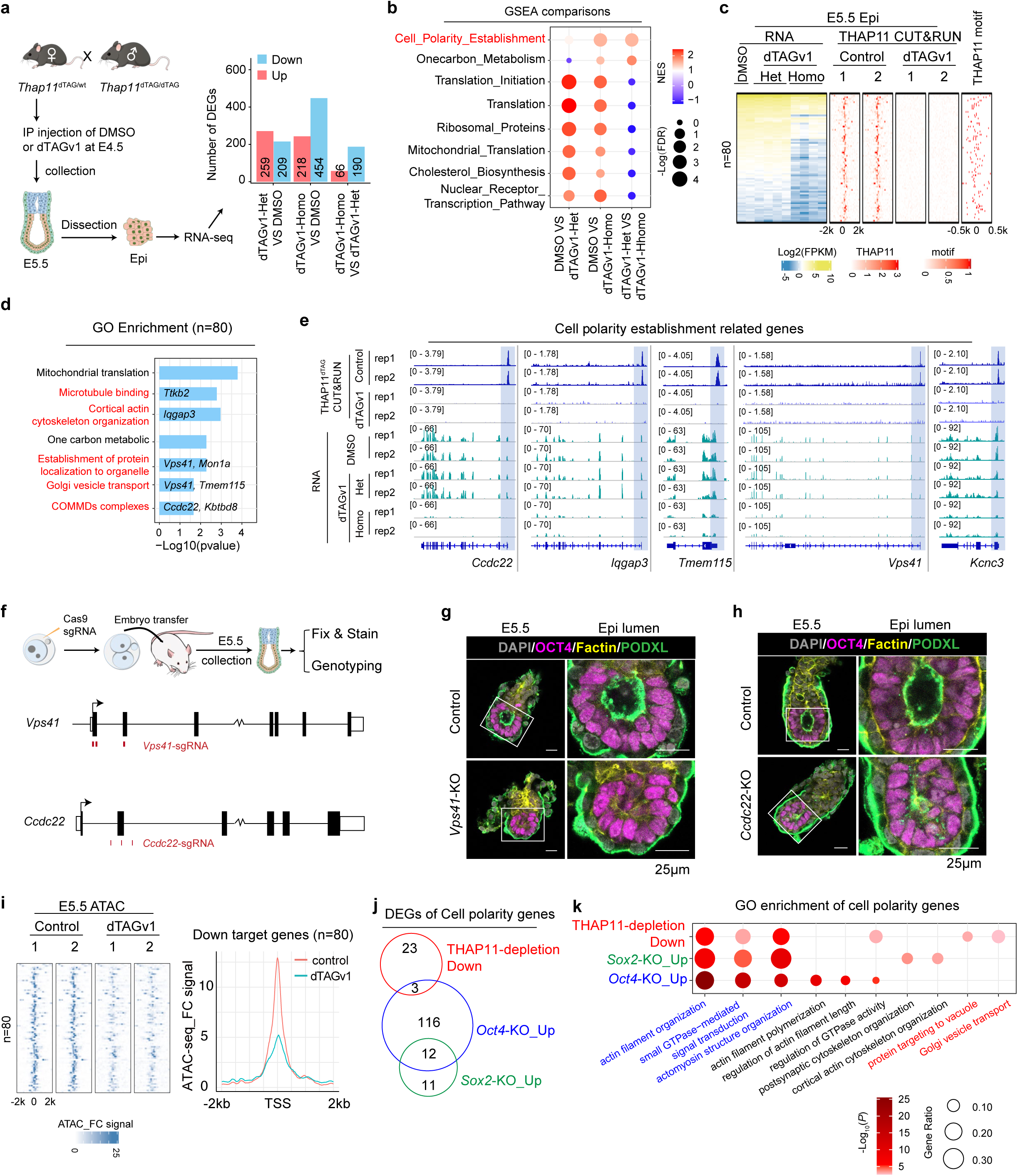
THAP11 regulates epiblast polarization through promoter binding. **a,** Schematic diagram of experimental design and the number of differentially expressed genes (DEGs) identified in each comparison. Fold change > 2, false discovery rate (FDR) < 0.05. **b,** Dotplots of GSEA comparisons among E5.5 DMSO, dTAGv1-Het, and dTAGv1-Homo Epi cells. **c,** Heatmaps showing the THAP11-bound DEGs in E5.5 Epi upon THAP11 loss, together with THAP11 motif occurrence profiles around their TSSs. **d,** Gene Ontology (GO) terms enriched for the THAP11-bound down-regulated genes (n=80) with examples. Red texts indicate terms related to cell polarity establishment. **e,** Genome browser view of THAP11 targeted polarity gene examples showing THAP11 CUT&RUN and RNA-seq in E5.5 Epi cells. **f,** Schematic diagram of experimental design. **g-h,** Representative immunostaining images of OCT4, F-actin, and PODXL in control, *Vps41*-KO, and *Ccdc22*-KO E5.5 embryos from Supplementary information, Fig. S4c and d\. A high-resolution view of the boxed regions is shown on right. Scale bar, 25 μm. **i,** Heatmaps and density plots showing the enrichment and average ATAC–seq signal fold change (ATAC_FC) in the promoter regions of THAP11 bound DEGs (n=80) upon THAP11 loss, respectively. **j,** Venn diagram showing the overlapped polarity genes between downregulated in E5.5 THAP11-depleted Epi cells and upregulated in E4.5 *Oct4*- or *Sox2*-KO Epi cells. **k,** Dotplots showing GO terms enriched for the polarity genes down-regulated upon THAP11-depletion and upregulated upon *Oct4*- or *Sox2*-KO.

### THAP11 activates Epi cell polarity genes through promoter binding

To investigate the mechanism, we performed comparative gene expression analysis and identified 454 downregulated and 218 upregulated genes in dTAGv1-Homo relative to DMSO, and 190 downregulated and 66 upregulated genes in dTAGv1-Homo relative to dTAGv1-Het in E5.5 Epi cells following THAP11 depletion (**Fig. 2a, Supplementary information, Fig. S2i and Supplementary Table 2**). To identify the DEGs directly affected by THAP11 depletion, we performed CUT&RUN in E5.5 Epi cells and found that 90.17% THAP11 peaks are mapped to the gene promoter regions (**Supplementary information, Fig. S3a-b**). Importantly, the THAP11 peaks are lost in the THAP11-depleted E5.5 Epi cells (**Supplementary information, Fig. S3c-d**), demonstrating the specificity of the binding peaks. Consistently, the THAP11 binding peaks are highly enriched for THAP11 binding motif (**Supplementary information, Fig. S3d-e**). Furthermore, integrative analysis revealed that 80 of the downregulated gene promoters are occupied by THAP11 (**Fig. 2c and Supplementary Table 2**). Since low-input CUT&RUN can inherently miss peaks leading to underestimation, we performed motif analysis on the 454 downregulated genes and found that 75.77% (344/454) contained at least one THAP11 motif in their promoter regions (**Supplementary information, Fig. S3f**), suggesting a direct role of THAP11 in regulating these genes. Gene Ontology (GO) analysis of these 80 direct targets, including *Iqgap3*, *Vps41*, and *Ccdc22*, revealed the enrichment in terms for microtubule binding, cortical actin cytoskeleton organization, and Golgi vesicle transport (**Fig. 2d-e**), which are essential for cell polarity establishment. These data collectively suggest that THAP11 directly regulate cell polarity genes through promoter binding during Epi morphogenesis.

### THAP11-activated *Vps41* and *Ccdc22* are vital for epiblast morphogenesis

To demonstrate that activation of key THAP11 target genes is essential for Epi morphogenesis, we focused on *Vps41* and *Ccdc22*. Previous studies have shown that loss of *Vps41* function results in gastrulation defects 23, and *Ccdc22* is essential for ITGB1 recycling 24. Notably, *Itgb1* knockout leads to Epi morphogenesis failure 25. Using a single-generation knockout strategy 26, we generated E5.5 knockout embryos for morphological analysis (**Fig. 2f**). Immunostaining revealed that both E5.5 *Vps41*-KO and *Ccdc22-*KO embryos exhibited reduced PODXL expression at the Epi lumen (**Fig. 2g-h and Supplementary information, Fig. S4a-d**), indicating impaired epiblast morphogenesis. Collectively, these data indicate that THAP11 may regulate epiblast morphogenesis, at least in part, through activating *Vps41* and *Ccdc22* expression.

To investigate whether THAP11 regulates target gene expression through affecting chromatin accessibility in E5.5 Epi, we performed ATAC-seq (**Supplementary information, Fig. S4e**). Comparative analysis revealed that THAP11 loss significantly reduced chromatin accessibility at the promoters of THAP11-bound downregulated genes (for example *Ccdc22*, *Iqgap3*, *Vps41*, and *Tmem115*) (**Fig. 2i and Supplementary information, Fig. S4f**). Taken together, these data indicate that THAP11 contributes to establishment of promoter chromatin accessibility, and that loss of THAP11 leads to defective polarity gene expression in E5.5 Epi cells.

### THAP11 regulates epiblast polarization independent of OCT4 and SOX2

A recent study showed that OCT4 and SOX2 function as repressors of Epi polarization 11 as their knockout results in pre-activation of polarity genes in E4.5 Epi cells. We thus asked whether THAP11 has some regulatory relationship with OCT4 and SOX2. We found that acute THAP11 degradation neither affected the expression of the pluripotency regulator *Otx2* nor *Oct4* or *Sox2* (**Supplementary information, Fig. S4g**), suggesting that THAP11 regulates Epi polarization independently of these known regulators. Additionally, neither *Oct4*-nor *Sox2*-knockout significantly affected *Thap11* expression in E4.5 Epi cell (**Supplementary information, Fig. S4h**), suggesting that the repressive role of OCT4 and SOX2 in Epi cells are independent of THAP11. Furthermore, an integrated analysis showed that only 3 out of the 26 THAP11-activated polarity genes overlapped with those of OCT4-repressed polarity genes (**Fig. 2j and Supplementary information, Fig. S4i, and Supplementary Table 3**). Nevertheless, GO analysis revealed that THAP11-, OCT4-, and SOX2-regulated polarity genes are both enriched for terms related to actin filament organization, GTPase signal transduction, and actomyosin structure organization (**Fig. 2k**), suggesting that they regulate different genes involved in similar processes. Notably, THAP11-regulated polarity genes are uniquely enriched for proteins targeting to vacuole and Golgi vesicle transport (**Fig. 2k**). Collectively, these data demonstrate that THAP11 regulates Epi polarization independent of OCT4 and SOX2, acting through a distinct transcriptional program.

### THAP11 is required for TE polarization and blastocyst formation

After demonstrating a role of THAP11 in Epi polarization, we next asked whether it also has a role in the first wave of embryonic polarization during 8-cell to morula development. Immunostaining revealed the presence of THAP11 from oocyte to blastocyst stage (**Fig. 3a**), consistent with a potential role in the first wave of embryonic polarization. Unexpectedly, THAP11 cannot be completely depleted by dTAG13 treatment in *Thap11*^dTAG/dTAG^ morulae (**Fig. 3b**), precluding the use of the *Thap11*^dTAG/dTAG^ model to investigate THAP11’s role in this wave of cell polarization. Since both the maternal deposited mRNA and ongoing transcription contribute to the THAP11 protein pool in preimplantation embryos, we reasoned that a combination of genetic knockout and dTAG13-mediated protein degradation might achieve complete depletion (**Supplementary information, Fig. S5a**). To this end, we first generated *Thap11*^wt/-^ mice by deleting the entire THAP11 coding region (**Supplementary information, Fig. S5b**). Consistent with previous reports ^19^, *Thap11*^-/-^ mice were embryonic lethal between E4.75 to E5.5 (**Supplementary information, Fig. S5c-e**), indicating successful knockout. We then generated *Thap11*^dTAG/-^ mouse model by crossing *Thap11*^wt/-^ with *Thap11*^dTAG/dTAG^ (**Supplementary information, Fig. S5f**). Notably, these *Thap11*^dTAG/-^ mice are fertile and developmentally normal, with an average litter-size of 10.22 pups based on 22 litters obtained from *Thap11*^dTAG/-^ and *Thap11*^dTAG/dTAG^ crosses (**Supplementary information, Fig. S5g**), indicating that the dTAG knock-in does not compromise THAP11 function under physiological conditions. Importantly, dTAG13 treatment resulted in complete depletion of THAP11 protein in *Thap11*^-/-^ morulae (**Fig. 3c and Supplementary information, Fig. S5f**). Together, these results support that the THAP11^dTAG/-^ mice can serve as a robust model for studying THAP11’s function during preimplantation development.

**Fig. 3.**
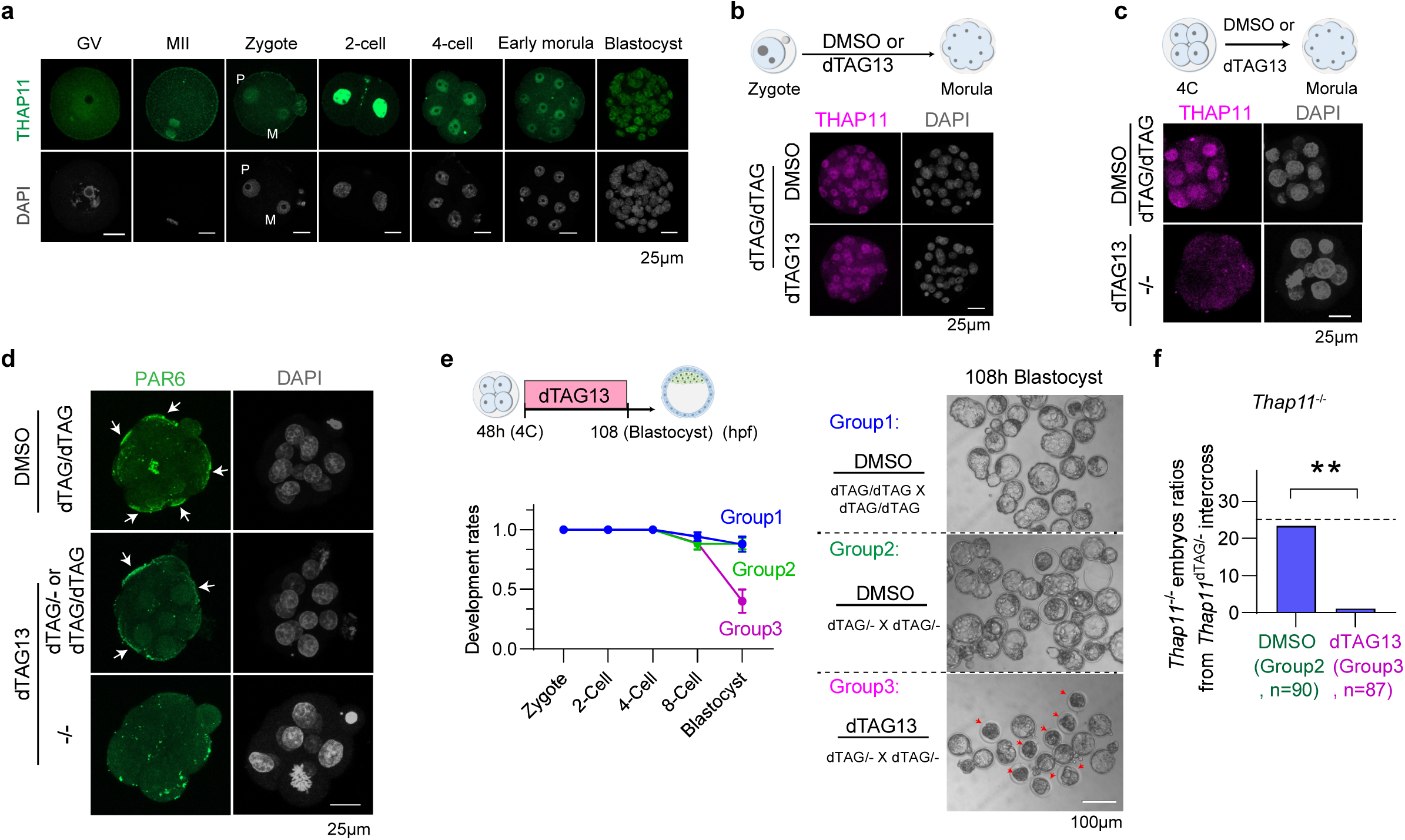
THAP11 is required for TE polarization and blastocyst formation. **a**, Immunostaining of THAP11 during mouse oocyte and early embryo development. Scale bar, 25 μm. **b,** Immunostaining of THAP11 in *Thap11*^dTAG/dTAG^ morulae with or without dTAG13 treatment. Scale bar, 25 μm. **c,** Immunostaining of THAP11 at morula of DMSO treated *Thap11*^dTAG/dTAG^ and dTAG13 treated *Thap11*^-/-^ embryos. Scale bar, 25 μm. **d,** Immunostaining of PAR6 at morula of DMSO treated *Thap11*^dTAG/dTAG^ and dTAG13 treated *Thap11*^-/-^ embryos. Three independent experiments are performed. Scale bar, 25 μm. **e,** Left panel: schematic diagram of the experimental design (top) and the development rate of embryos from group1 (DMSO treated *Thap11*^dTAG/dTAG^ embryos), group2 (DMSO treated embryos from *Thap11*^dTAG/-^ intercrosses), and group3 (dTAG13 treated embryos from *Thap11*^dTAG/-^ intercrosses) (bottom). The data are presented as mean ± SEM. Right panel: representative images of blastocyst stage embryos from different group (group1, n = 74; group2, n = 55, group3, n = 58; n represents the total number of embryos from two independent experiments). Scale bar, 100 μm. Red arrows indicate representative arrested embryos in group3. Scale bar, 100 μm. **f,** Genotype ratios of *Thap11*^-/-^ blastocysts from *Thap11*^dTAG/-^ intercrosses with or without dTAG13 treatment (group2, n=90; group3, n=87 blastocysts; four independent experiments. Chi-squared test, **p<0.01.

To directly test THAP11’s function during TE polarization, we treated embryos with dTAG13 for 24h starting from 4-cell embryo and collected at early morula (EM) stage for immunostaining. The results showed that apical PAR6 staining was diminished in the THAP11-depleted (dTAG13-*Thap11*^-/-^) embryos (**Fig. 3d and Supplementary information, Fig. S5h**), suggesting a role of THAP11 in the first wave of embryo polarization. Since embryo polarization from 8-cell to morula is vital for blastocyst formation ^4,27,28^, we next sought to examine the developmental potential of the THAP11-depleted embryos (**Fig. 3e**). Expectedly, over 90% embryos from group1 (DMSO treated *Thap11*^dTAG/dTAG^) and group2 (DMSO treated *Thap11*^dTAG/dTAG^, *Thap11*^dTAG/-^, and *Thap11*^-/-^) developed to blastocysts, whereas about 60% of embryos from group3 (dTAG13 treated *Thap11*^dTAG/dTAG^, *Thap11*^dTAG/-^, and *Thap11*^-/-^) failed to reach blastocyst stage (**Fig. 3e**). Importantly, genotyping of blastocysts from group2 revealed a 23% ratio of *Thap11*^-/-^ blastocysts (**Fig. 3f**), consistent with the expected mendelian ratio (25%). In contrast, almost no THAP11-depleted (dTAG13 treated *Thap11*^-/-^) group3 embryos reached blastocyst (**Fig. 3f**). Additionally, we observed that THAP11-depleted embryos had a comparable total cell number to control embryos at early morula stage (**Supplementary information, Fig. S5i**) but exhibited a marked reduction in cell number by 108h, when control embryos had reached the blastocyst stage, accompanied by extensive DNA fragmentation (**Supplementary information, Fig. S5j**), suggesting increased cell death. Consistently, extensive cell death has also been reported in the Pard6b-knockdown model ^29^, raising the possibility that polarity deficiency may contribute to increased cell death. Together, these results demonstrate a critical role of THAP11 in the first wave of embryo polarization and blastocyst formation.

### THAP11 activates TE polarity genes through promoter binding

To understand how THAP11 regulates TE polarization, we performed RNA-seq for early morula embryos (**Supplementary information, Fig. S6a**) and confirmed successful *Thap11* knockout (**Supplementary information, Fig. S6b**). Comparative analysis revealed 1,702 downregulated and 1,300 upregulated genes in response to THAP11-depletion (**Fig. 4a and Supplementary Table 4**). GSEA comparisons revealed that cell polarity establishment and cell junction organization were significantly disrupted after THAP11-depletion (**Fig. 4b**). Importantly, THAP11 depletion did not significantly affect the expression of morula regulator *Nr5a2* ^30–32^ or embryo polarity regulators *Rhoa*, *Tfap2c*, and *Tead4* ^10^ (**Supplementary information, Fig. S6c**), suggesting that THAP11 regulates this wave of embryo polarization independently of these known regulators. Consistent with previous notion that embryo polarity establishment is required for TE fate specification ^4,10^, THAP11-depletion resulted in a significant decrease in the expression of TE marker genes *Gata3* and *Cdx2* (**Supplementary information, Fig. S6c**).

**Fig. 4.**
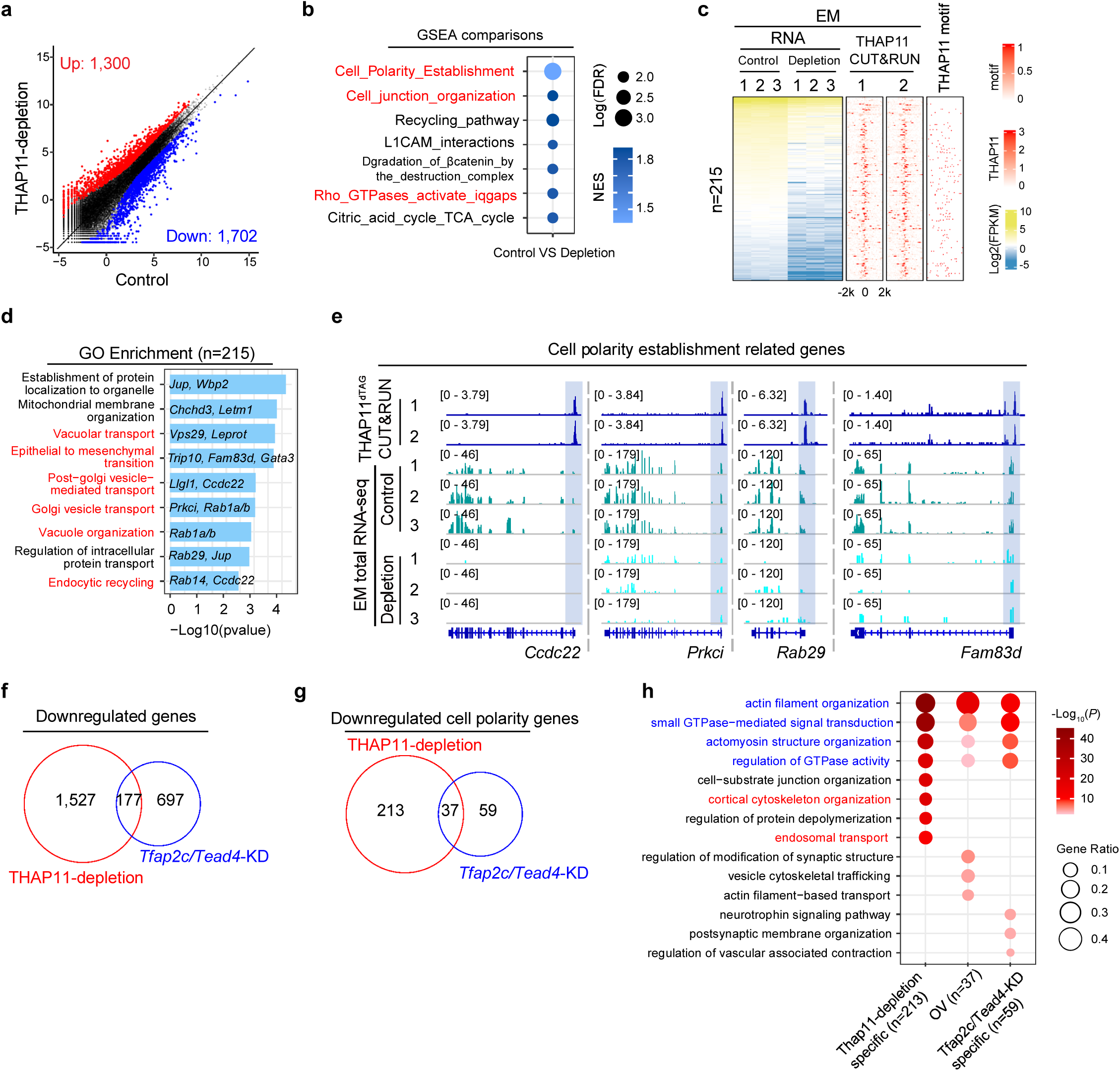
THAP11 regulates TE polarization through promoter binding. **a,** Scatter plots comparing gene expression profiles of early morula (EM) with or without dTAG13 treatment. The x and y axis of the dot plots are Log_2_CPM from RNA-seq. Fold change > 2, FDR < 0.05. **b,** Dotplots of GSEA comparisons between control and THAP11-depleted EM. **c,** Heatmaps showing the THAP11-bound DEGs in EM upon THAP11 loss, together with THAP11 motif occurrence profiles around their TSSs. **d,** Gene Ontology (GO) terms enriched for the THAP11-bound down-regulated genes (n=215) with examples. **e,** Genome browser view of THAP11targeted polarity gene examples showing THAP11 CUT&RUN and RNA-seq in EM. **f,** Venn diagram showing the overlapped downregulated genes between THAP11-depletion and *Tfap2c*/*Tead4*-KO in EM. **g,** Venn diagram showing the overlapped downregulated polarity genes between THAP11-depletion and *Tfap2c*/*Tead4*-KO in EM. **h,** Dotplots showing GO terms enriched for the polarity genes down-regulated upon THAP11-depletion and *Tfap2c*/*Tead4*-KO in EM.

To identify the genes directly affected by THAP11 depletion, we performed CUT&RUN in early morula embryos (**Supplementary information, Fig. S6d**). We found that THAP11 peaks are mainly localized to promoter regions that are also enriched for the THAP11 motif, but not for the NR5A2, TFAP2C, or TEAD4 motifs (**Supplementary information, Fig. S6f-g**). Integrative analyses of CUT&RUN and RNA-seq data revealed 215 downregulated genes with promoter THAP11 binding (**Fig. 4c and Supplementary Table 4**). Since low-input CUT&RUN data can underestimate binding events, we performed motif analysis of the 1,487 down-regulated genes without detectable promoter CUT&RUN signal. Notably, 69.5% (1,034/1,487) of these genes contained at least one THAP11 motif in their promoter regions (**Supplementary information, Fig. S6h**), supporting a direct role for THAP11-promoter binding in their regulation. GO analysis of the 215 downregulated direct targets (including *Ccdc22* and *Prkci*) revealed enrichment of terms for vacuolar transport, Golgi vesicle transport, and endocytic recycling (**Fig. 4d-e and Supplementary information, Fig. S6i**), processes essential for cell polarity establishment. Collectively, these data suggest that THAP11 regulates cell polarity genes by directly binding to their promoters during the first wave of embryo polarization.

Previous studies have revealed a role for TFAP2C and TEAD4 on this wave of polarization ^10^. We thus explored a possible relationship between THAP11 and these known TFs. We reanalyzed the TFAP2C/TEAD4-KD RNA-seq datasets ^10^. Comparative analysis of the affected genes revealed that the majority of the THAP11 downregulated genes were not affected by TFAP2C/TEAD4-KD (**Fig. 4f**), consistent with THAP11 regulates polarization independently of TFAP2C/TEAD4 (**Supplementary information, Fig. S6c**). Moreover, 85.2% (213 out of 250) of the THAP11-regulated polarity genes, including *Ccdc22*, *Llgl1*, and *Prkci*, were not affected by TFAP2C/TEAD4-KD (**Fig. 4g and Supplementary Table 5**). Interestingly, GO analysis does reveal shared enrichment of actin filament organization and actomyosin structure organization among the THAP11- and TFAP2C/TEAD4-regulated genes (**Fig. 4h**). Intriguingly, THAP11-specific down-regulated polarity genes are uniquely enriched for cortical cytoskeleton organization and endosomal transport (**Fig. 4h**). Taken together, these findings suggest that the first wave of embryo polarization is co-regulated by THAP11 and TFAP2C/TEAD4, with largely independent and partially convergent transcriptional programs.

### THAP11 overexpression promotes polarity gene expression and premature clustering of apical proteins

To help clarify whether THAP11 functions as an instructive inducer or as a permissive regulator of cell polarization, we microinjected *Thap11* mRNA into one of the two-cell blastomere and cultured them to early 8-cell for immunostaining of the polarity marker PAR6 **(Supplementary information, Fig. S7a)**. The results showed a significant increase in large PAR6 clusters in THAP11 overexpressed early 8-cell blastomeres compared with the control blastomeres **(Supplementary information, Fig. S7b)**, suggesting that THAP11 overexpression promotes premature clustering of apical proteins PAR6. Previous Studies have shown that the first wave of cell polarization involves sequential clustering of apical proteins, followed by expansion and centralization of the apical domain ^33^. Notably, overexpression of TFAP2C/TEAD4 can advance the formation of apical protrusions enriched in apical proteins by approximately one cell cycle, whereas activated RhoA promotes subsequent apical-domain expansion ^33^. Consistent with this model, our data suggest that THAP11 overexpression is sufficient to promote premature apical-protein clustering, indicative of an earlier onset of apical polarization. However, THAP11 overexpression alone did not induce expansion of the apical domain, suggesting that additional factors, such as RhoA-dependent signaling, may be required for subsequent apical-domain maturation and expansion. Intriguingly, we also found that nuclear PAR6 enrichment significantly increased upon THAP11-OE **(Supplementary information, Fig. S7c)**, although nuclear PAR6 function remains unknown.

To understand how THAP11-OE promote the clustering of PAR6, we performed RNA-seq of THAP11-OE E8C embryos (**Supplementary information, Fig. S7d**) and confirmed the increase in *Thap11* expression (**Supplementary information, Fig. S7e**). Comparative transcriptomic analysis revealed 347 upregulated and 177 downregulated genes in response to THAP11-OE in E8C embryos (**Supplementary information, Fig. S7f and Supplementary Table 6**). GO analysis of upregulated genes revealed significant enrichment of terms in epithelium morphogenesis, vacuolar localization, and myotube differentiation in response to THAP11-OE (**Supplementary information, Fig. S7g**). Specifically, GO analysis of upregulated polarity genes revealed that GTPase-mediated signal transduction, actin filament organization, and vesicle cycle were significantly increased after THAP11-OE (**Supplementary information, Fig. S7h**), which might explain the clustering of PAR6. Taken together, our *Thap11* overexpression data suggest that THAP11 most likely functions as an instructive inducer of cell polarization.

### THAP11 is required for PrE polarization and maturation

At E3.5, the ICM cells consist of intermixed Epi- and PrE-progenitors that heterogeneously express lineage marker genes and progressively differentiate into naïve Epi cells and polarized, cavity-facing, and mature PrE cells ^34,35^. This second lineage segregation is initially guided by a transcription regulatory network involving OCT4, GATA6, and fibroblast growth factor (FGF) signaling ^35–38^, and is subsequently matured and stabilized by cell polarization ^5,6^. Although PKC signaling and GTPase RAC1 have been shown to be required for PrE polarization ^5,6^, a transcription factor involved in regulating this process has not been identified. Given our findings that THAP11 is required for both 8-cell to morula TE polarization and E4.5-E5.5 Epi polarization, we next investigated whether and how THAP11 may regulate E3.5-E4.5 PrE polarization. To this end, we first tested the degradation efficiency and found that THAP11 cannot be completely depleted by dTAG13 treatment in the *Thap11*^dTAG/dTAG^ blastocysts (**Supplementary information, Fig. S8a**). We therefore employed THAP11^dTAG/-^ mouse model and achieved efficient THAP11 depletion in the *Thap11*^-/-^ blastocysts following dTAG13 treatment from early to late blastocyst stage (LB) (**Fig. 5a-b**). Notably, THAP11-depletion did not dramatically alter the number of PrE cells (GATA6^+^ cells) (**Fig. 5c**), indicating that THAP11 is dispensable for PrE cell fate initiation.

**Fig. 5.**
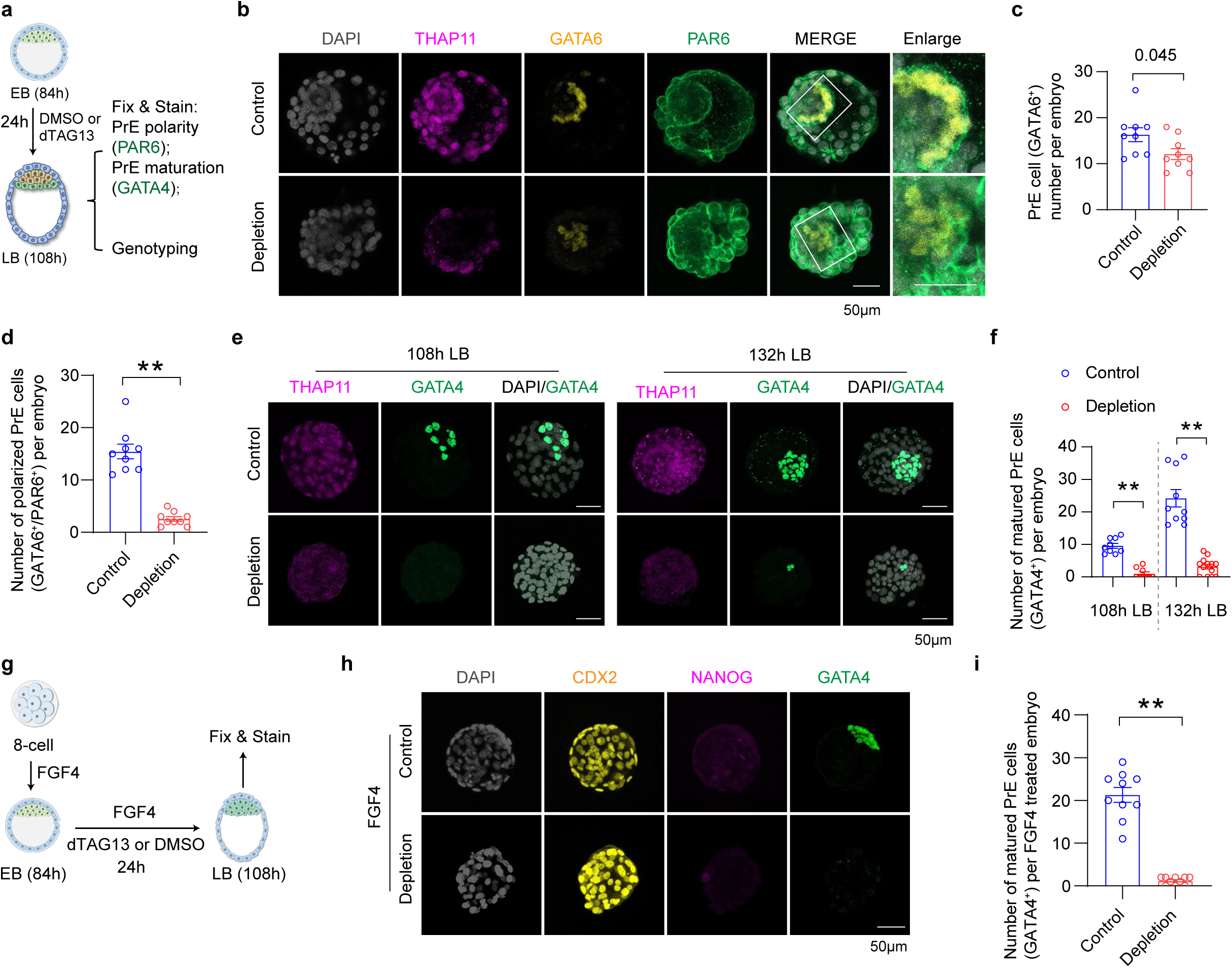
THAP11 depletion causes PrE polarization and maturation defects. **a,** Schematic diagram of experimental design. **b,** Z-stack projection of THAP11, GATA6, and PAR6 immunostaining in control and THAP11-depleted LB. An enlarged view of the boxed regions is shown on right. Individual Z-stack optical sections are showed in supplementary information, Fig. S8b. **c,** Number of PrE cells (GATA6^+^) per embryos of control (n=9) and THAP11-depletion (n=9). **d,** Number of polarized PrE cells per embryo of control (n=9) and THAP11-depletion (n=9). **e,** Immunostaining of THAP11 and GATA4 in control and THAP11-depleted LB. **f,** Number of mature PrE cells (GATA4^+^) per LB of control (108h, n= 9; 132h, n=10) and THAP11-depletion (108h, n= 9; 132h, n=13) at the indicated time. **g,** Schematic diagram of experimental design. **h,** Immunostaining of lineage markers CDX2, NANOG, and GATA4 in FGF4 treated control and THAP11-depleted LB. **i,** Number of mature PrE cells (GATA4^+^) of FGF4 treated control (n=10) and THAP11-depletion (n=10) LB. Scale bar, 50 μm. n represents the total number of embryos from two independent experiments; Quantitative data are shown as mean ± SEM; ∗∗p < 0.01, Student’s t test.

Importantly, in control embryos (DMSO-*Thap11*^dTAG/dTAG^), PrE cells were properly aligned along the cavity side and exhibited apical PAR6 expression, whereas the majority of THAP11-depleted (dTAG13-*Thap11*^-/-^) PrE cells failed to establish apical PAR6 expression (**Fig. 5b, 5d and Supplementary information, Fig. S8b-c**), demonstrating a critical role of THAP11 in PrE polarization. As cell polarization is essential for PrE maturation ^5,6^, we next investigated whether THAP11-depletion disrupts PrE maturation (**Fig. 5a**). To this end, we stained the mature PrE marker GATA4 which revealed that the THAP11-depleted blastocysts failed to specify mature PrE cells, even after extended culture for 48h (**Fig. 5e-f**), indicating a failure of PrE maturation. Intriguingly, zygotic THAP11 knockout embryos can develop to E4.5 (**Supplementary information, Fig. S8d**) and showed no significant change in Epi cell numbers, but exhibited a moderate decrease in mature PrE cells (**Supplementary information, Fig. S8e-g**), suggesting that both maternal and zygotic THAP11 contribute to efficient PrE maturation.

Previous studies has showed that luminal deposition of FGF4 secreted by Epi cells plays a critical role in initiation, positioning, and maturation of PrE cells ^39,40^. To exclude that the disruption of PrE maturation in THAP11-depleted blastocysts are secondary effects from potential defects in Epi cells, we converted all the ICM cells into PrE by treating embryos with FGF4 starting from 8-cell (**Fig. 5g**), which is a well-established method for Epi/PrE segregation studies ^6,35,37^. Consistent with previous studies ^35^, immunostaining of cell lineage markers revealed that no Epi cells (NANOG^+^), but only TE (CDX2^+^) and mature PrE cells (GATA4^+^), were found in the FGF4 treated control LB (**Fig. 5h and Supplementary information, Fig. S8h**), suggesting that all the ICM cells have been successfully converted to PrE cells. In contrast, no mature PrE cells were found in the FGF4 treated THAP11-depleted LB (**Fig. 5h-i**), suggesting that the intrinsic defects in PrE cells is the cause of PrE maturation failure upon THAP11-depletion. Collectively, these data support that THAP11 is required for PrE polarization and maturation.

### THAP11 regulates PrE polarization through promoter binding

To understand how THAP11 regulates PrE polarization, we performed RNA-seq. To capture the earlier molecular changes that cause defective PrE polarization, and to avoid confounding from potential Epi defects, we converted all ICM to PrE cells by FGF4 treatment and collected PrE cells at the mid-blastocyst stage after 16h dTAG13 treatment (**Supplementary information, Fig. S9a**). After confirming successful *Thap11* knockout (**Supplementary information, Fig. S9b**), we performed comparative transcriptomic analysis which revealed 615 upregulated and 1,015 downregulated genes in response to THAP11-depletion in PrE cells (**Fig. 6a and Supplementary Table 7**). GO analysis and GSEA comparisons both revealed that actin filament organization, cell junction localization, and myosin complex were significantly disrupted after THAP11-depletion (**Fig. 6b and Supplementary information, Fig. S9c**).

**Fig. 6.**
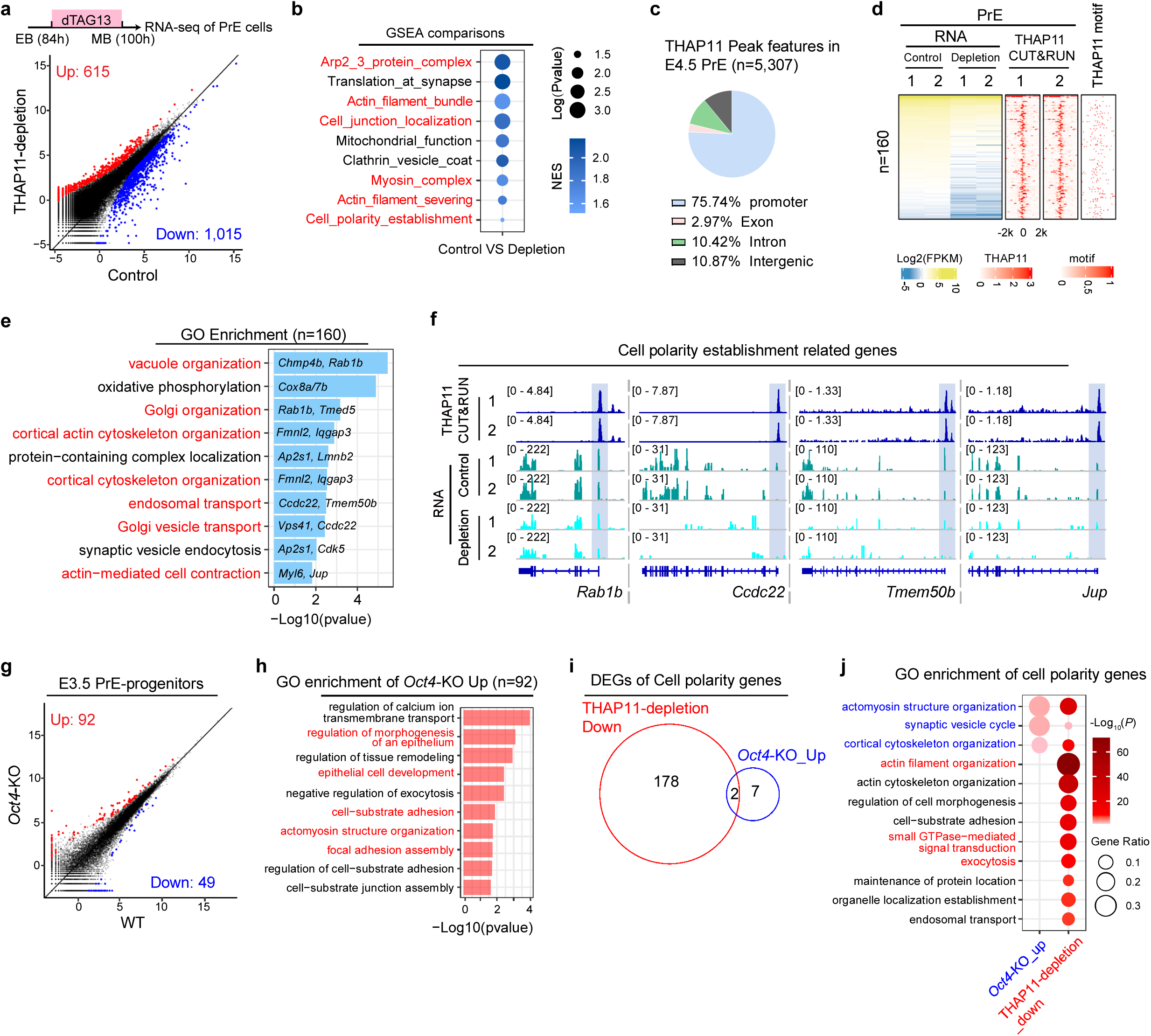
THAP11 regulates PrE polarization through promoter binding. **a,** Upper panel: schematic diagram indicates dTAG13 treatment window and sample collection time. Lower panel: scatter plots comparing the gene expression profiles of 100h PrE cells with or without dTAG13 treatment. The x and y axis of the dot plots are Log_2_CPM from RNA-seq. Fold change > 2, FDR < 0.05. **b,** Dotplots of GSEA comparisons between control and THAP11-depleted PrE cells. **c,** The genomic distribution of THAP11 binding peaks based on THAP11 CUT&RUN data in E4.5 PrE cells. **d,** Heatmaps showing the THAP11-bound DEGs in PrE cells upon THAP11 loss, together with THAP11 motif occurrence profiles around their TSSs. **e,** Gene Ontology (GO) terms enriched for THAP11-bound down-regulated genes (n=160) with examples. **f,** Genome browser view of THAP11 targeted polarity gene examples showing THAP11 CUT&RUN and RNA-seq in PrE cells. **g,** Scatter plots comparing the gene expression profiles of E3.5 PrE progenitors upon *Oct4*-KO. The x and y axis of the dot plots are Log_2_CPM from RNA-seq. Fold change > 2, FDR < 0.05. **h,** Representative GO terms enriched in the upregulated genes (n=92) in response to *Oct4*-KO. **i,** Venn diagram showing the overlapped polarity genes between downregulated in THAP11-depleted PrE cells and upregulated in *Oct4*-KO PrE progenitors. **j,** Dotplots showing GO terms enriched for the polarity genes that are down-regulated upon THAP11-depletion and upregulated upon *Oct4*-KO.

To identify genes directly affected by THAP11 depletion, we performed CUT&RUN of PrE cells (**Supplementary information, Fig. S9d**). We found that THAP11 peaks are mainly mapped to promoter regions where the THAP11 binding motif is highly enriched, but not that of the OCT4 or SOX2 binding motifs (**Fig. 6c and Supplementary information, Fig. S9e-f**). Integrative analyses of CUT&RUN and RNA-seq data revealed 160 downregulated genes with promoter THAP11 binding (**Fig. 6d and Supplementary Table 7**). Since low-input CUT&RUN tend to underestimate binding events, we performed motif analysis of the 855 downregulated genes without detectable promoter THAP11 CUT&RUN signal and identified 65.7% (562/855) of these genes contained at least one THAP11 binding motif in their promoter regions (**Supplementary information, Fig. S9g**), supporting a direct role for THAP11-promoter binding in their regulation. GO analysis revealed that the 160 downregulated direct targets, including *Rab1b* and *Ccdc22*, are enriched for vacuole organization, Golgi organization, and cortical actin cytoskeleton organization (**Fig. 6e-f**), processes critical for cell polarization. Collectively, these data suggest that THAP11 directly regulates cell polarity genes through promoter binding during PrE polarization.

### THAP11 activates PrE polarization program independently of OCT4

Since knockout OCT4 and SOX2 results in premature activation of cell polarity in E3.5 ICM cells 11, it is possible that OCT4 and SOX2 might also repress PrE polarity genes. To test this possibility, we analyzed public single-cell RNA-seq datasets of SOX2-KO 41 and OCT4-KO 42 E3.5 ICM cells. Transcriptome comparisons of E3.5 PrE-progenitors (with higher *Gata6* and *Pdgfra* expression than that of the Epi progenitors) revealed 92 upregulated and 49 downregulated genes in response to OCT4-KO, whereas almost no DEGs in response to SOX2-KO (**Fig. 6g, Supplementary information, Fig. S9h, and Supplementary Table 8**), suggesting that only OCT4 but not SOX2 may serve as a repressor of PrE polarization. Importantly, GO analysis revealed that the 92 upregulated genes are enriched for terms related to morphogenesis of epithelial cells, epithelial cell development, cell adhesion, and actomyosin structure organization (**Fig. 6h**), consistent with previous study showing OCT4-KO leads to premature activation of polarity in E3.5 ICM cells. In summary, these findings suggest that OCT4 serves as a repressor of PrE polarization.

Since THAP11 and OCT4 respectively activates and represses PrE polarity genes, we asked whether they have common gene targets. Comparative analysis of their regulated genes revealed that only 2 genes are overlapped between THAP11-depletion downregulated genes and OCT4-KO upregulated genes (**Supplementary information, Fig. S9i**), indicating their independent function. Importantly, only 2 out of the 180 THAP11-activated polarity genes are repressed by OCT4 in PrE (**Fig. 6i and Supplementary Table 9**). Intriguingly, GO analysis revealed that THAP11- and OCT4-regulated polarity genes are both enriched for the terms of actomyosin structure organization and cortical cytoskeleton organization (**Fig. 6j**). Notably, THAP11-regulated polarity genes are uniquely enriched for terms related to actin filament organization, GTPase signal transduction, and exocytosis (**Fig. 6j**). Collectively, these data demonstrate that THAP11 regulates PrE polarization independent of OCT4, and functions through a distinct transcriptional program.

### THAP11 regulates three waves of polarization through common and stage-specific chromatin binding

Having demonstrated that THAP11 regulates all three waves of cell polarization, independently of the known regulators, we next sought to identify common and stage-specific mechanisms by which THAP11 regulates these polarization events. By comparing THAP11 binding profiles in early morula (EM) embryos, E4.5 PrE, and E5.5 Epi, we identified 5 clusters (C1-C5) with stage-specific binding features and a large cluster (C6) with conserved binding features (**Fig. 7a and Supplementary information, Fig. S10a-b**). Intriguingly, only EM-specific peaks (C1) are predominantly located at distal regions, whereas other stage-specific peaks (C2-C5) are mainly located at promoter regions (**Fig. 7b**), underscoring the distinct THAP11-chromatin binding preferences during different polarization events. To further characterize these stage-specific THAP11 binding features, we performed integrative analyses of CUT&RUN and RNA-seq datasets, and identified stage-specific binding associated downregulated genes (**Supplementary information, Fig. S10c and Supplementary Table 10**). GO analysis of these stage-specific downregulated genes revealed that they are enriched for terms related to cell polarization, such as “regulation of microtubule cytoskeleton” in C1, “establishment of cell polarity” in C2, “actin filament organization” in C3, “maintenance of protein location” in C4, and “myotube differentiation” in C5 (**Fig. 7c**), suggesting conserved functions in regulating cell polarization despite different aspects of cell polarization.

**Fig. 7.**
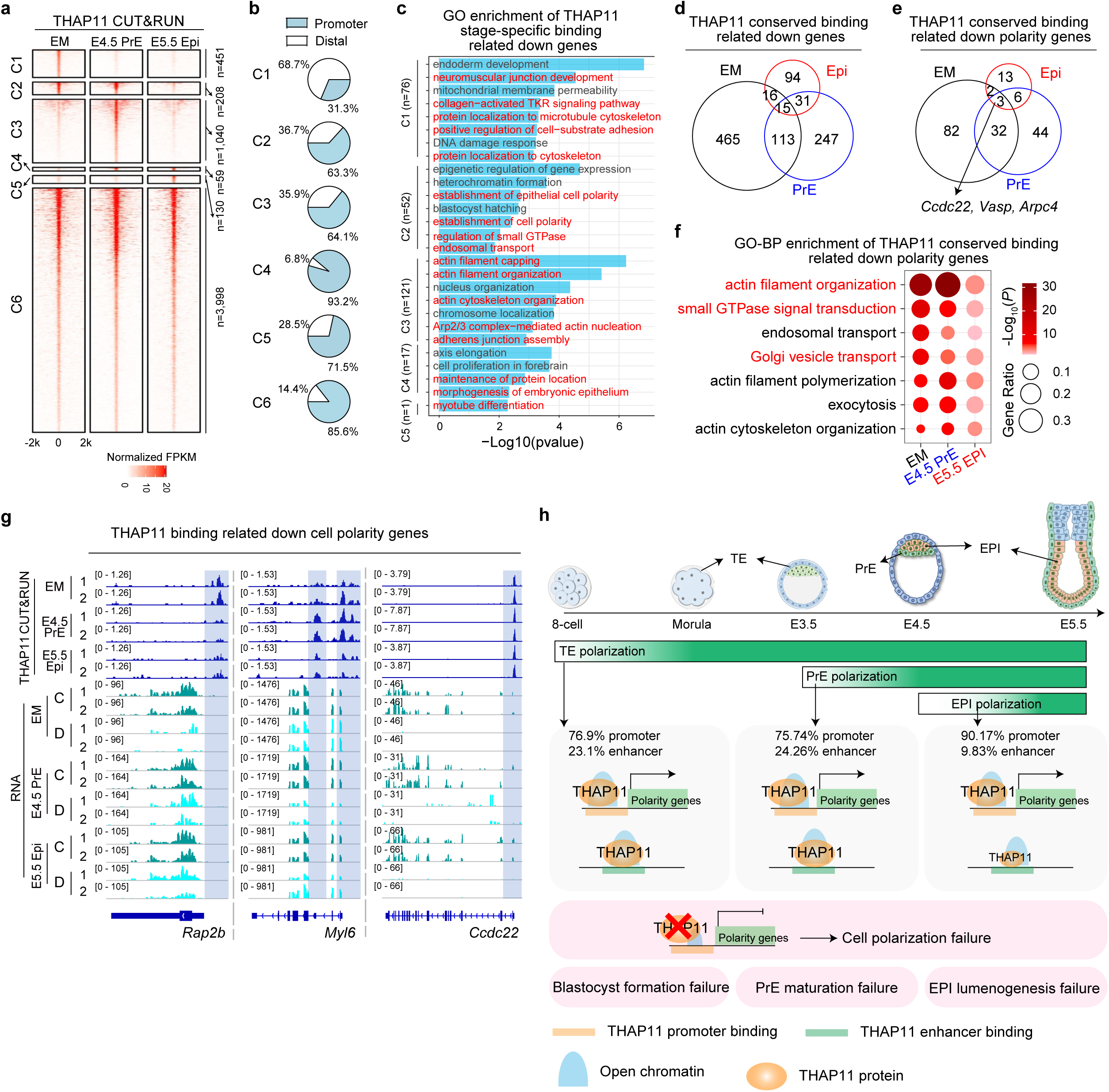
THAP11 regulates polarity programs by promoter and distal chromatin binding. **a,** Heatmaps showing the dynamics of THAP11 chromatin binding profile in EM, E4.5 PrE, and E5.5 Epi at promoter and distal regions. **b,** Pie chart showing the percentage of promoter and distal occupancy. **c,** Representative GO terms enriched in downregulated genes (FDR<0.05, LogFC<0) associated with each stage-specific binding cluster (C1-C5). Red texts indicate terms related to cell polarity establishment. **d,** Venn diagram showing the overlaps among THAP11 conserved binding related downregulated genes (FDR<0.05, LogFC<0) in EM (n=609), E4.5 PrE (n=406), and E5.5 Epi (n=156). **e,** Venn diagram showing the overlaps among THAP11 conserved binding related downregulated polarity genes in EM (n=119), E4.5 PrE (n=85), and E5.5 Epi (n=23). **f,** Representative GO-BP terms enriched among THAP11 conserved binding-related downregulated polarity genes at the indicated stages. **g,** Genome browser view of THAP11 binding related down-regulated polarity genes upon THAP11 depletion showing CUT&RUN and RNA-seq at indicated stages. **h,** Diagram illustrating the role of THAP11 in the three waves of polarization during early embryogenesis. Across three sequential cell polarization events, THAP11 activates polarity genes predominantly through promoter binding. Loss of THAP11 disrupts all three waves of cell polarization, leading to blastocyst formation failure, impaired PrE maturation, and defective Epi lumenogenesis, respectively.

In addition to the stage-specific binding, we also identified 3,998 common THAP11 binding peaks predominantly (85.6%) located at promoter regions (**Fig. 7a-b and Supplementary information, Fig. S10a-b**). Integrative analysis of the CUT&RUN and RNA-seq dataset revealed 981 downregulated genes are associated with these common THAP11 binding peaks (**Supplementary information, Fig. S10d and Supplementary Table 11**). GO analysis of these 981 genes revealed enrichment in terms related to endosomal transport, vesicle organization, and Golgi vesicle transport (**Supplementary information, Fig. S10e**). Among the 981 genes, 465 are specifically downregulated in EM, 247 in E4.5 PrE, and 94 in E5.5 Epi upon THAP11-depletion (**Fig. 7d and Supplementary information, Fig. S10d**). Focusing on cell polarity genes, we identified 182 genes with conserved THAP11 binding and are downregulated in different stages with THAP11 depletion, including 82 EM-specific, 44 E4.5 PrE-specific, and 13 E5.5 Epi-specific downregulated polarity genes (**Fig. 7e**). GO analysis revealed that these stage-specific downregulated genes are commonly associated with Biological Processes, such as actin filament organization, GTPase signal transduction, and Golgi vesicle transport (**Fig. 7f**); Cellular Components, such as apical junction complex and Golgi subcompartment; and Molecular Functions, such as GTPase activity and actin binding (**Supplementary information, Fig. S10f**).

Taken together, these results suggest that THAP11 regulates three waves of cell polarization through a combination of divergent and conserved chromatin binding (**Fig. 7g-h**). Although different sets of polarity genes are regulated by THAP11 binding at different stages, all of them function in core polarity programs, such as actin filament organization, GTPase signal transduction, and Golgi vesicle transport, underscoring a fundamental requirement for THAP11 in cell polarization.

Integrating our findings with previous reports ^33,43^, we propose a conceptual framework for the sequential regulation of cell polarization in which THAP11 functions as a general regulator across all three waves of polarization, acting cooperatively with, or independently of, stage-specific regulators to coordinate these sequential polarization events. To further explore the potential basis of this context specificity, we performed motif enrichment analyses of THAP11 binding at different stages and identified candidate cofactors that may cooperate with THAP11 in different developmental contexts (**Supplementary table 12**). For example, KLF5/17/4 and GATA3 may help establish THAP11 function in EM (**Supplementary information, Fig. S10g**); KLF6/9 and GATA2/4/6 may contribute to THAP11 function in E4.5 PrE (**Supplementary information, Fig. S10h**); ETV1/2/4, MAZ, and TFDP1 may contribute to THAP11 function in E5.5 Epi (**Supplementary information, Fig. S10i**). Further work is needed to test whether any of the predicted factors indeed work with THAP11 to confer context specificity.

## Discussion

The importance of cell polarization in early mammalian embryonic development is evident from its requirement for promoting lineage segregation and increasing embryonic structural and functional complexity. Here, we demonstrate that THAP11 plays an important role in activating polarity genes in all three waves of cell polarization during early embryogenesis. Phenotypically, stage-specific depletion of THAP11 disrupts all three waves of cell polarization and leads to the failure of blastocyst formation, PrE maturation, and Epi lumenogenesis, respectively (**Fig. 7h**). Mechanistically, THAP11 regulates the three waves of polarization through conserved and stage-specific chromatin binding. Intriguingly, both stage-specific and conserved THAP11-chromatin bindings regulate core polarity programs, including actin filament organization, GTPase signal transduction, and Golgi vesicle transport. Together, our study demonstrates THAP11 is a general regulator of cell polarity establishment during early embryogenesis.

Intriguingly, besides the disruption of polarity in E5.5 Epi cells upon THAP11 loss, we also observed the reduction of polarity marker PAR6 in the endoderm cells surrounding the epiblast (**Fig. 1**). This raises the possibility that loss of polarity of the endoderm disrupts polarization of the epiblast in a non-cell autonomous way. However, several lines of evidence support a direct role for THAP11 in epiblast polarization. First, THAP11 directly regulates essential polarity genes, including *Vps41* and *Ccdc22*, in epiblast cells. Second, genetic disruption of these target genes phenocopies the epiblast morphogenesis defects. Third, acute THAP11 depletion in our 3D mESC self-organization model disrupts epithelial polarization and lumenogenesis in the absence of extraembryonic tissues. Together, these findings support a cell-autonomous requirement for THAP11 in epiblast epithelialization, although not excluding the possibility that non-cell-autonomous mechanisms also contribute *in vivo*. The functional contribution of endoderm polarity to epiblast morphogenesis remains to be further investigated.

In mammalian embryogenesis, cell polarization is a major driver of morphogenesis, emerging sequentially in TE cells at early morula stage, in PrE cells at E4.5, and in Epi cells at E5.5. Despite its central role, how these sequential polarization events are spatial-temporally regulated remains poorly understood. In the past few years, great progress has been made in identifying TFs that regulate cell polarization in a stage-specific manner, including the activators TFAP2C and TEAD4 in TE polarization ^10^, and the repressors OCT4 and SOX2 in Epi polarization ^11,12^. However, these stage-specific cell polarization regulators cannot explain how cell polarization is sequentially regulated during early embryogenesis. Given that cell polarization is a highly conserved and organized process, we propose the existence of a general transcriptional activator that governs all three waves of cell polarization, acting in concert with stage-specific regulators to coordinate these sequential polarization events. Here, we demonstrate that THAP11 is such a general transcriptional activator, providing a conceptual framework for the sequential regulation of cell polarization (**Fig. 7h**). During 8-cell to morula TE polarization, THAP11 and TFAP2C/TEAD4 independently activate polarity genes, predominantly through promoter binding (**Fig. 3-4**) and enhancer binding ^10,44^, respectively. During E3.5 to E4.5 PrE polarization, THAP11 acts alongside the repressor OCT4 to regulate polarity genes predominantly through promoter binding (**Fig. 5-6**) and enhancer binding ^45^, respectively. Similarly, during E4.5 to E5.5 Epi polarization, THAP11 cooperates with the repressors OCT4/SOX2, independently regulating polarity genes mainly through promoter binding (**Fig. 1-2**) and enhancer binding ^11,41^, respectively. Consistent with this model, core polarization programs, including actomyosin structure organization and cytoskeleton organization, are co-regulated by these factors across developmental stages (**Fig. 2, 4, and 6**). Notably, THAP11 uniquely regulates a subset of polarity genes involved in Golgi vesicle transport, including *Vps41* and *Ccdc22* (**Fig. 2, 4, and 6**), highlighting a conserved transcriptional module that links vesicle trafficking to polarity establishment across preimplantation to peri-implantation development. Although we identified candidate cofactors, including KLF5/17/4 and GATA3 in EM, KLF6/9 and GATA2/4/6 in E4.5 PrE, and ETV1/2/4, MAZ, and TFDP1 in E5.5 Epi, that may contribute to the stage-specific activity of THAP11, functional validation of these factors and the underlying mechanisms remain to be investigated.

Our findings reveal that both common and stage-specific THAP11 bindings exert conserved functions in polarity regulation. As expected, stage-specific bindings regulate distinct aspects of cell polarity (**Fig. 7c**). In contrast, common THAP11 bindings exhibit unexpected plasticity, regulating different sets of polarity genes across developmental stages **(Fig. 7e)**. This dynamic regulatory landscape suggests a model where stage-specific co-regulators may compensate for THAP11’s loss in regulating specific subset of polarity genes. Notably, these genes are commonly enriched for the core polarity programs, including actin filament organization, GTPase signal transduction, and Golgi vesicle transport **(Fig. 7f)**, highlighting a fundamental requirement for THAP11 in regulating cell polarity establishment. Our findings establish THAP11 as a central regulator of cell polarization and raise the question of whether increasing THAP11 activity is sufficient to enhance or advance polarity programs, or even drive *de novo* polarization. Our preliminary data indicate that THAP11 overexpression promotes premature apical-protein clustering at the E8C stage and enhances the expression of core polarity programs, including GTPase-mediated signal transduction, actin filament organization, and vesicle cycling. Whether increased THAP11 activity can similarly promote or advance polarization during the PrE and EPI waves remains to be determined. Although we have identified THAP11-targeted polarity genes at different developmental stages, the specific target genes that mediate the polarization defects caused by THAP11 depletion remain to be determined. Notably, we demonstrated that *Vps41* and *Ccdc22* are essential THAP11 target genes in E5.5 Epi cells, as their depletion impaired Epi polarization. However, the detailed mechanisms by which VPS41 and CCDC22 contribute to Epi polarization remain to be elucidated.

THAP11 has an established function in mitochondria gene regulation ^46–48^, consistent with our observation that a subset of downregulated targets is associated with mitochondrial function (**Fig. 2d**). Mitochondria plays an important role in hepatocytes polarization ^49^, raising the possibility of an indirect THAP11-mitochondria-polarization regulatory axis. However, mitochondria depletion during preimplantation development does not disrupt blastocyst formation ^50^, indicating that TE polarization can proceed normally under these conditions. This suggests that mitochondrial function is unlikely to play a major role in early embryonic polarization. Moreover, our acute degradation strategy and short time treatment window likely minimize the emergence of secondary metabolic defects, supporting a direct role of THAP11 in regulating polarity gene expression, which is consistent with its direct binding to these gene promoters. Since both THAP11 and cell polarization are evolutionarily conserved ^8,51,52^, functional demonstration of its evolutionarily conserved roles in cell polarization in other species await to be shown.

## Resource availability

Further information and requests for resources and reagents should be directed to the lead contact, Yi Zhang

## Data and code availability

Public data used in this study are listed in Supplemental Table 16. Source data are provided with this paper. This paper does not report original code.

## Materials and Methods

### *Thap11*-KO and *Thap11*-dTAG mouse generation

All animal experiments were performed in accordance with the protocols of the Institutional Animal Care and Use Committee at Harvard Medical School. All mice were kept under specific pathogen-free conditions within an environment controlled for temperature (20-22°C) and humidity (40-70%), and were subjected to a 12 h light/dark cycle. All mice had a C57BL/6J genetic background. Primers used for genotyping are listed in **Supplementary Table 13**.

*Thap11*-dTAG knock-in mice were generated following a previous protocol with modifications17. Briefly, 2-cell embryo (20hpf) were injected with *Thap11* donor DNA (30 ng/μl), Cas9 protein (25 ng/μl) and sgRNA (50 ng/μl each) using a Piezo impact-driven micromanipulator (Primer Tech, Ibaraki, Japan). After 2 h KSOM incubation, 2-cell embryos were transferred into oviducts of pseudo-pregnant ICR strain mothers (Charles River). F0 chimera mice were backcrossed with wild-type C57BL/6J mice for at least two generations. Genotyping was performed with mouse tail lysed in lysis buffer (50 mM Tirs-HCl, 0.5% Triton, 400 μg/ml Proteinase K) at 55°C overnight. For F0 and F1 mice genotyping, the primers outside the homology arm were used. For F2 and beyond mice genotyping, inner primers were used. The primers are included in **Supplementary Table 13**.

*Thap11*-KO heterozygous mice were generated following a similar procedure. Zygotes were injected with Cas9 protein (25 ng/μl) and two *Thap11*-sgRNAs (50 ng/μl each). After one day KSOM incubation, 2-cell embryos were transferred into oviducts of pseudo-pregnant ICR strain mothers. F0 chimera mice were backcrossed with wild-type C57BL/6J mice for at least two generations. KO genotyping primers and sgRNA sequence are listed in **Supplementary Table 13**. *Thap11*^dTAG/-^ mice were generated by crossing *Thap11*-KO heterozygotes with *Thap11*-dTAG mice.

Droplet Digital PCR (ddPCR) was used for detecting the copy number of *Thap11*-dTAG knock-in allele in F1 mice. Briefly, 250 ng purified DNA template was digested by incubation with Haelll Enzyme (NEB) at 37°C for 1h, and then inactivated at 80°C for 5 mins. A final 30 ng DNA was used as template for PCR. *Fkbp* was used for knock-in detecting, *mRPP30* was used as internal control of two copy genome. The PCR primers are listed in **Supplementary Table 13**. The mice with single *Fkbp* copy were used for subsequent mating (**Supplementary Table 14**).

### Preparation and collection of mouse oocytes and embryos

All experiments involved in oocytes and embryo preparation were performed as previously described 17,53, with minor modifications. Female mice (7-8 weeks) were super-ovulated through an initial injection of 5 IU pregnant mare serum gonadotropin (PMSG, BioVendor, RP1782725000), followed by a 5 IU injection of human chorionic gonadotropin (hCG, Sigma, C1063) 48 hours later. Oocyte-cumulus complexes were collected 14 hours post hCG injection. Sperms were collected from the cauda epididymis of adult male mice (8-12 weeks). The sperm suspension was capacitated for 1 hour in 200 μl HTF medium (Millipore, MR-070-D). Oocytes were then incubated with spermatozoa for 6-hours. The time when sperms were added to oocytes was considered as 0 hpf. Two-nuclear zygotes were cultured in the KSOM medium (Millipore, MR-106-D) under a humidified atmosphere of 5% CO_2_ at 37°C for their development.

### dTAG13 or dTAGv1 treatment

dTAG13 (Tocris, 6605) or dTAGv1 (Tocris, 6914) was reconstituted in DMSO to a 50 mM stock. For THAP11 degradation in mESCs, dTAG13 was diluted in mESCs culture medium to 0.5 μM. For THAP11 degradation in preimplantation embryos, dTAG13 was diluted in KSOM to 0.5 μM. Embryos were washed with KSOM with dTAG13 for at least three times, then cultured in KSOM with dTAG13 for further development. For THAP11 degradation in post-implantation embryos *in vivo*, intraperitoneal (IP) injections were employed. 1 mg dTAGv-1 was initially dissolved in 25 μl DMSO and subsequently diluted with 475 μl 10% castor oil (Sigma, C5135) to prepare the injection solution. Mice (∼28 g each) were injected with 1 mg dTAGv1 every 12 hours to maintain sustained THAP11 degradation.

### Cell culture and establishment of *Thap11*^dTAG^ mESC cell line

Undifferentiated E14 mESCs were cultured on 0.1% gelatin coated plates with 2i and LIF condition. E14 mESCs were grown in DMEM (Gibco, 11960069), supplemented with 15% fetal bovine serum (Sigma-Aldrich, F6178), 2 mM GlutaMAX (Gibco, 35050061), 1 mM sodium pyruvate (Gibco, 11360), 1x MEM NEAA (Gibco, 11140050), 0.084 mM 2-mercaptoethanol (Gibco, 21985023), 1 mM sodium pyruvate (Gibco, 11360070), 100 U/ml penicillin-streptomycin (Gibco, 15140122), 1000IU/ml LIF (Millipore, ESG1107), 0.5 μM PD0325901 (Tocris, 4192), and 3 μM CHIR99021 (Tocris, 4423). To establish *Thap11^dTAG^* mESC cell line, *Thap11* donor DNA and *Thap11*-sgRNA inserted pX330.puro (addgene, #110403) plasmids were co-transfected with Lipofectamine™ 2000 (Invitrogen, 11668030). Cells were then selected with puromycin (Gibco, A1113803), and the single cell-derived colonies were then picked for genotyping and further analysis. Genotyping primers are listed in **Supplementary Table 13**. Cells were cultured in an incubator containing 5% CO_2_ at 37 °C.

### Western blot

Cells were lysed in 1 x LDS buffer (Invitrogen, NP0007) at 1 × 10^4^ cell/μL final concentration and heated at 100 °C for 10 min. Samples were run on NuPAGE 4-12% gel (Invitrogen, NP0322BOX) and transferred onto PVDF Transfer Membrane. Primary antibodies used include anti-THAP11 (1:250, MAB5727, R&D systems), anti-HA (1:1000, 3724S, Cell Signaling Technology), and anti-GAPDH (1:50000, 60004-1-Ig, Proteintech). Protein bands were detected with ECL kit (Thermo Fisher Scientific, 32209) and imaged by iBright 1500.

### Immunofluorescence

Oocytes or embryos were fixed in 4% paraformaldehyde at room temperature 20min or 4 overnight and permeabilized in 0.5% Triton X-100 - 0.1% Polyvinylalcohol-PBS (PBST-PVA) at room temperature for 1 h. After blocking with 1% BSA in PBST-PVA, the samples were incubated in primary antibodies diluted in PBST-PVA containing 1% BSA overnight at 4 . Antibodies were used as follows: anti-THAP11 (1:300, MAB5727, R&D systems), anti-HA (1:200, 3724S or 2367S, Cell Signaling Technology), anti-OCT4 (1:200, sc-5279, Santa Cruz), anti-OCT4 (1:500, ab181557, Abcam), anti-GATA6 (1:300, 5851T, Cell Signaling Technology), anti-PODXL (1:300, MAB1556-SP, R&D systems), anti-PAR6 (1:300, sc-166405, Santa Cruz), anti-GATA4 (1:200, R&D Systems, MAB2606-SP), anti-NANOG (1:200, Abcam, ab80892), anti-CDX2 (1:500, R&D Systems, AF3665-SP), anti-CDX2 (1:200, MU392A-5UC, BioGenex). Following three washes with PBST-PVA, the embryos were incubated with appropriate secondary antibodies for 1 hour at room temperature: Donkey anti-Mouse IgG-Alexa Fluor 568 (1:500, Invitrogen, A10037), Donkey anti Rabbit IgG (H+L)-Alexa Fluor 488 (1:500, Invitrogen, A-21206), Donkey anti-Goat IgG (H+L)-Alexa Fluor 647 (1:500, Invitrogen, A-21447), Phalloidin-iFluor 647 Reagent (1:1000, ab176759, Abcam), or Phalloidin-iFluor 555 Reagent (1:1000, ab176756, Abcam). Hoechst 33342 (10 μg/ml, Sigma) was used for DNA staining. The fluorescence signals were imaged using a confocal laser scanning microscope (Zeiss, LSM800). The fluorescent intensity was quantified using ImageJ. In Fig. 5, cells were classified as CDX2-, GATA4/6-, or NANOG-positive when the corresponding nuclear fluorescence signal was clearly above the background level and cells were classified as PAR6-positive when the corresponding apical fluorescence signal was clearly above the cytoplasmic level, whereas cells lacking a distinguishable signal above background were classified as negative.

### Immunosurgery

ICMs were isolated as previously described 17. Briefly, blastocysts at E3.5 or E4.5 stages were collected by removing the zona pellucida with Acidic Tyrode’s solution (Millipore). Embryos were then treated with anti-mouse serum antibody (Sigma-Aldrich, M5774-2ML, 1:5 dilution in KSOM) for 30 min at 37 °C. After washing three times with KSOM, embryos were treated with guinea pig complement (Millipore, 1:5 dilution in KSOM) for another 20 min at 37 °C. Then, the TE cells were removed by a glass pipette. As described previously 17, to obtain PrE cells, we treated E2.5 embryo with 500 ng/mL FGF4 and 1 μg/mL heparin to induce PrE cell fate in all ICM cells as embryos reach blastocyst stage, followed by performing immunosurgery to isolate the PrE population.

### CUT&RUN, ATAC and RNA-seq library preparation and sequencing

CUT&RUN assays were performed as previously described with minor modifications 17. For mESCs CUT&RUN with more than 1 × 10^4^ cells, cells were resuspended in 50 μl washing buffer (20 mM HEPES/pH=7.5, 150 mM NaCl, 0.5 mM spermidine and 1× protease inhibitor) with activated Concanavalin A Magnetic Beads (Polysciences, 86057-3) for 10 mins at room temperature (RT), then samples were incubated with anti-HA (1:100, ab9110, lot#GR3437862-1, Abcam) overnight at 4°C. Samples were then incubated with 2.8 ng/μl pA-MNase (home-made) for 2 hrs at 4 °C. Subsequently, samples were incubated with 200 μl pre-cooled 2 μM CaCl2 for 20 mins at 4 °C and quenched by the addition of 23 μl 10×stop buffer (1.7 M NaCl, 20 mM EGTA, 100 mM EDTA, 0.02% Digitonin, 250 µg/ml glycogen and 250 µg/ml RNase A). DNA fragments were released by incubation at 37°C for 15 mins. Then, 2.5 μl 10% SDS and 2.5 μl 20 mg/ml Protease K (Thermo Fisher) were added and incubated at 55 °C for at least 1 h. DNA was extracted by phenol-chloroform followed by ethanol precipitation. For low input mESCs and embryo CUT&RUN, some modifications were made. Briefly, mESCs or zona-free embryos were resuspended in 50 μl wash buffer with activated Concanavalin A Magnetic Beads for 10 mins at RT, then samples were incubated with anti-HA overnight at 4°C. The subsequent procedures are the same as those for mESCs described above. Sequencing libraries were prepared with the NEBNext Ultra II DNA library preparation kit for Illumina (New England Biolabs, E7645S).

ATAC-seq was performed as previously described with some modifications 17,53,54. Briefly, isolated oocytes were digested with adapter-loaded Tn5 for 15 mins at 37°C, and stopped by stop buffer (100 mM Tris/pH=8.0, 100 mM NaCl, 40 µg/ml Proteinase K and 0.4% SDS) and incubated overnight at 55°C. Then, 5 µl of 25% Tween-20 was added to quench SDS. Sequencing libraries were prepared with NEBNext High-Fidelity 2×PCR Master Mix (NEB, M0541S). RNA-seq library was prepared with SMART-Seq® Stranded Kit (Takara Bio, 63444) using 5,000 mESCs. All libraries were sequenced by NextSeq 550 system (Illumina) with paired-ended 75-bp reads (**Supplementary Table 15**).

### RNA-seq data processing and analysis

Raw sequencing reads were trimmed using Trimmomatic 55 (v0.39) to remove sequencing adaptors, and subsequently mapped to the GRCm38 genome using STAR 56 (v2.7.8a). Gene expression was quantified with featureCounts 57 (v2.0.1) by counting reads mapped to each gene. Then, edgeR 58 (v3.32.1) was employed for normalization and differential expression analysis. Genes with RPKM lower than 1 were defined as lowly expressed genes and excluded from the differential expression analysis. Differentially expressed genes (DEGs) were identified using likelihood ratio test (glmFit and glmLRT functions from edgeR). Genes with a false discovery rate (FDR) below 0.05 and an absolute value of fold change greater than 2 were defined as DEGs. Gene Ontology (GO) enrichment was performed using the R package clusterProfiler 59. Gene Set Enrichment Analysis (GSEA) was performed using clusterProfiler and enrichplot.

For analysis of E3.5 PrE progenitors in both *Oct4*-KO and *Sox2*-mzKO datasets, we first downloaded raw data from GSE159030 and GSE203194, respectively. Count matrices were generated for each sample and analyzed using Seurat (v5.3.0) 60. Briefly, data were normalized and scaled using default parameters. Cells with high expression of *Gata6* and *Pdgfra* were defined as PrE progenitors. In the Sox2-mzKO dataset, 21 PrE progenitor cells were identified (control: 9; mzKO: 12), while in the Oct4-KO dataset, 41 cells were detected (control: 30; KO: 11). We then generated two pseudo-bulk replicates for each condition and performed differential gene expression analysis.

### Analysis of cell polarity genes

To obtain all the cell polarity-related genes, we collected genes annotated with “Rho/GTPase”, “cell polarity”, or “actin cytoskeleton” in GENEONTOLOGY (https://www.geneontology.org/). After deduplicates, we obtained 2,285 cell polarity-related genes in **Supplementary Table 1**. The expressed genes were defined as FPKM ≥ 1.

### CUT&RUN data analysis

Raw sequencing reads were trimmed using Trimmomatic 55 (v0.39) to remove sequencing adaptors, and subsequently aligned to the GRCm38 reference genome using bowtie2 61 (v2.4.2) with parameters: --local --very-sensitive-local --no-unal --no-mixed --no-discordant --dovetail -I 10 -X 700 --soft-clipped-unmapped-tlen. PCR duplicates were removed by Picard MarkDuplicates 62 (v2.23.4) and reads with a mapping quality below 30 were removed. The mapped reads were further filtered to only retain proper paired reads with fragment length between 10 and 120 bp. Then, MACS2 63 (v2.2.7.1) was used to call significant peaks with parameters “-f BAMPE -B --SPMR -q 0.05 -g mm --keep-dup all”. To obtain highly conserved THAP11 binding sites, only peaks present in both biological replicates were defined as binding sites. The signal tracks were generated with deeptools 64 bamCoverage (v3.5.1) with bin size of 1 and normalized using CPM (counts per million). For z-score-normalized signal tracks, we first used bamCoverage with bin size of 100 to generate FPKM signals, then used a customized script to calculate the z score of each bin. Peak annotation was performed using the ChIPseeker 65 R package. Peaks located within ±2,500 bp of transcription start sites (TSS) were classified as promoter peaks, whereas peaks outside this region were considered distal peaks. The heatmaps of binding profiles were calculated with deeptools 64 computeMatrix (v3.5.1) using bigwig signal tracks as input and bin size of 10 and visualized in R with profileplyr and EnrichedHeatmap 66 packages.

### ATAC–seq data analysis

ATAC–seq data were processed with the ENCODE ATAC–seq pipeline using default parameters (v2.1.2, https://github.com/ENCODE-DCC/atac-seq-pipeline). Peaks located within ±2,500 bp of transcription start sites (TSS) were classified as promoter peaks, whereas peaks outside this region were considered distal peaks.

### Motif enrichment analysis

Motif enrichment analysis was performed using HOMER 67 (v4.11) findMotifsGenome.pl with the mm10 reference and parameter -size 200, using peak regions as input. Briefly, HOMER identifies over-represented motifs in the input peak regions by comparing them to matched random background sequences. Enrichment P values for all motifs in all peaks were then corrected for multiple testing by calculating adjusted P value using the Benjamini-Hochberg method. Motifs with an adjusted p-value less than 0.05 were considered significantly enriched.

### Motif occurrence analysis

Motif occurrence analysis was performed using HOMER 67 (v4.11) annotatePeaks.pl with parameters mm10 -size -500,500 -hist 20 -ghist for regions around gene TSSs. Motif files were downloaded from the JASPAR database 68 and manually converted to HOMER motif format. The JASPAR motif ID for TFs analyzed in this study THAP11 (MA1573.2), NR5A2 (MA0505.1), TFAP2C (MA0524.2) and TEAD4 (MA0809.2), and a log odds detection threshold of 6.0 was applied. The motif occurrence matrix was visualized in R with the EnrichedHeatmap 66 package.

## Acknowledgements

We thank members of the Zhang lab and Qingji Lyu for discussion during the study; Chengjie Zhou, Yota Hagihara, and Jun Wu for commenting on the manuscript. Meng Wang for initial bioinformatic analysis and helpful comments. This project was supported by NIH (R01HD116750) and the HHMI. YZ is an investigator of the Howard Hughes Medical Institute.

## Author contributions

Y.Z. conceived, supervised, and supported the project. Q-Y.Y. conceived the project; established the THAP11-dTAG mice, *Thap11*-KO mice and THAP11-dTAG mESCs; performed most experiments including RNA-seq, CUT&RUN, ATAC-seq etc. B-Y.W. performed mouse breeding, embryo preparation and collection; performed mESC 3D culture experiments, Thap11-OE experiments, Thap11-degradation experiments, ATAC-seq and libraries preparation; made major contribution during revision process. S.J. performed all the bioinformatic analysis. Q-Y.Y., S.J., B-Y.W., and Y.Z. wrote and revised the manuscript. All authors interpreted the data.

## Competing interests

Authors declare that they have no competing interests.

## Supplementary Materials

Supplementary information, Fig. 1 to 10; Supplementary Table 1 to 16 (separated files)

## Notes

### Competing Interest Statement

The authors have declared no competing interest.

