## Supplementary information, Fig. S1 to 10 for "The transcription factor THAP11 is a general regulator of early embryonic polarization"

- 1
- 2
- 3
- 4
- 5
- 6
- 7
- 8
- 9
- 10
- 11
- 12
- 13

- 2
- 3
- 4
- 5

6
78
9
10
11
12
13

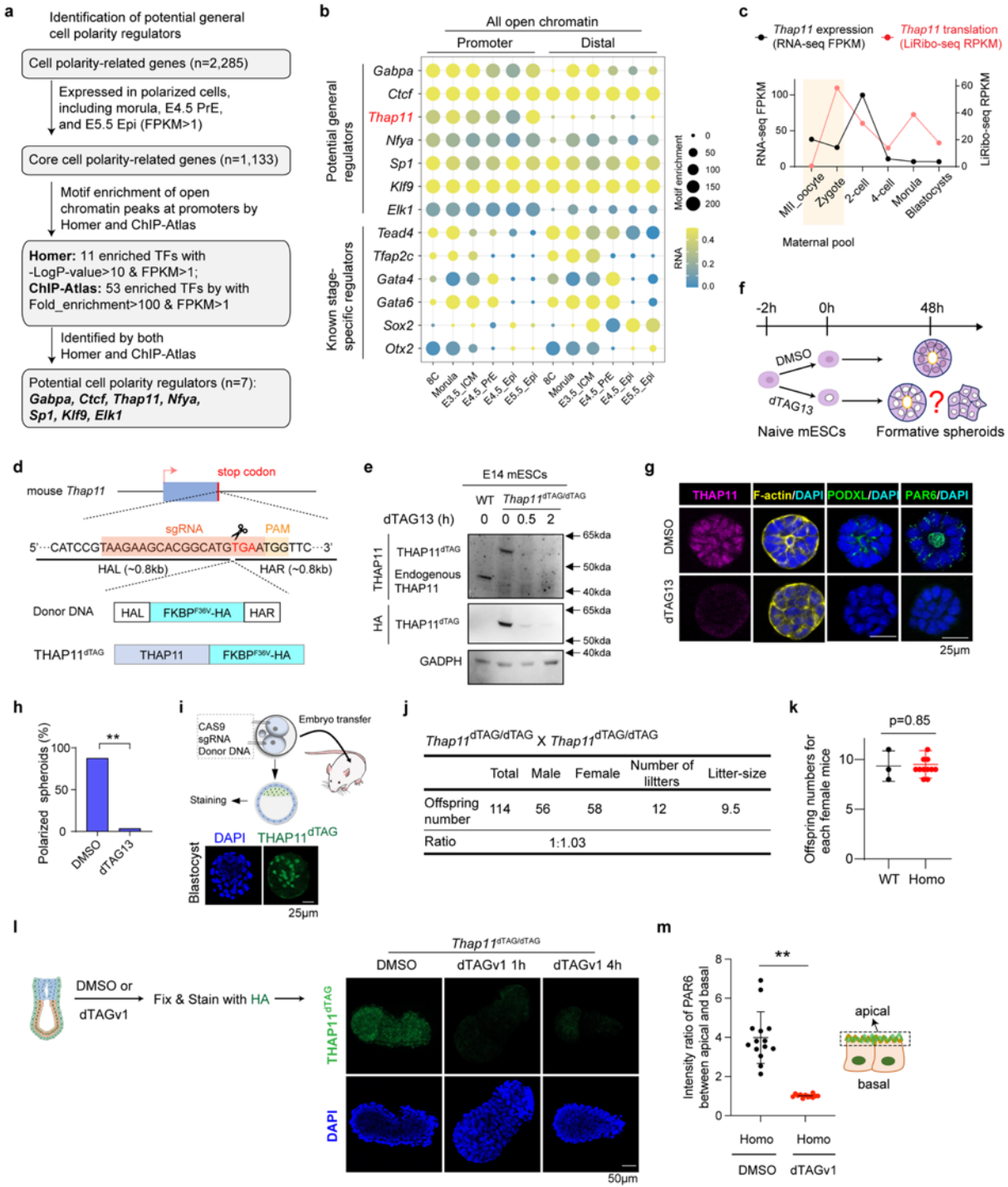

**Supplementary information, Fig. S1. Identifying THAP11 as a potential regulator of cell polarization and generation of THAP11-dTAG mouse line.** **a**, Schematic illustration for identifying candidate regulators underlying the three waves of cell polarization during early embryogenesis. **b**, Enrichment of TF motifs at promoter and distal ATAC-seq peaks of the

indicated stages. **c**, The dynamics of *Thap11* expression at transcription (RNA-seq, FPKM) and translation (LiRibo-seq<sup>1</sup>, RPKM) in mouse oocytes and early embryos. **d**, Diagram illustration of HA-FKBP (F36V) knock-in at the 3' end of *Thap11* gene and the THAP11<sup>dTAG</sup> fusion protein. **e**, Western blot analysis confirming THAP11<sup>dTAG</sup> knock-in and dTAG13-mediated THAP11<sup>dTAG</sup> degradation in mES cells. The blots were incubated with anti-THAP11 and anti-HA, respectively. GAPDH was used as a loading control. **f**, Schematic illustration of the experimental design. **g**, Immunostaining of 48h 3D cultured mESCs with DMSO or dTAG13 treatment. Scale bar, 25 μm. **h**, The ratio of polarized spheroids. Chi-squared test, \*\*p<0.01. **i**, Diagram illustration for *Thap11*<sup>dTAG/dTAG</sup> mouse generation and HA staining confirming the dTAG knock-in cells in blastocysts. Scale bar, 25 μm. **j**, Statistics of pup numbers from THAP11<sup>dTAG/dTAG</sup> × THAP11<sup>dTAG/dTAG</sup> crosses, indicating normal function of the THAP11<sup>dTAG</sup> fusion protein. **k**, Comparisons of pup numbers. Mean ± SEM; p-value, Student's t test. **l**, Left panel: experimental design. Right panel: Immunostaining of Homo embryos with or without dTAG13 treatment. **m**, Intensity ratio of PAR6 between apical and basal part in E5.5 Homo Epi with (n=12) or without (n=14) dTAGv1 treatment. \*\*p < 0.01, Student's t test.

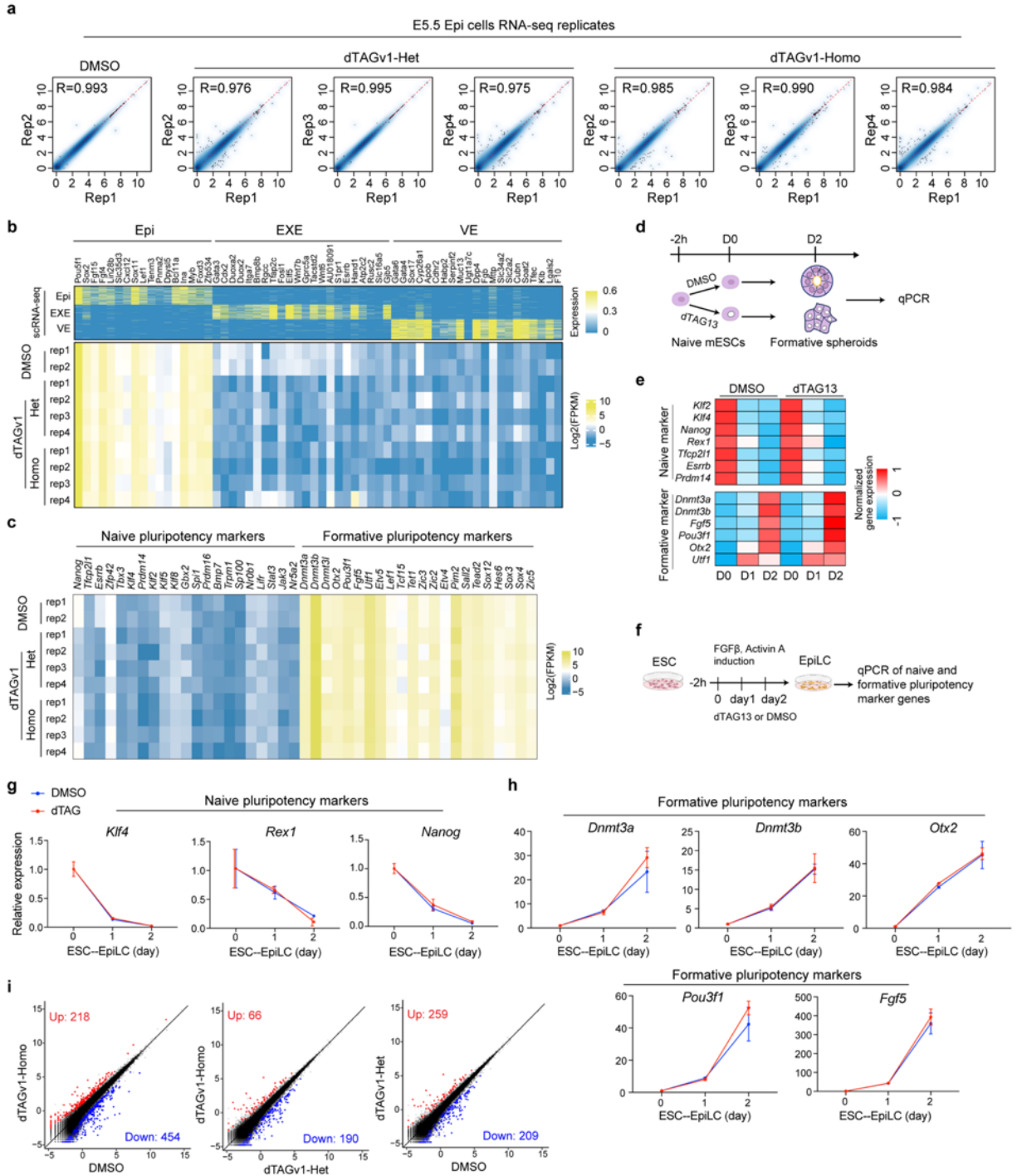

**Supplementary information, Fig. S2. THAP11 does not regulate pluripotency transition in epiblast cells.** **a**, Correlation of replicates for total RNA-seq in DMSO-*Thap11*<sup>dTAG/dTAG</sup> (DMSO-Homo), dTAGv1-*Thap11*<sup>dTAG/wt</sup> (dTAGv1-Het), and dTAGv1-*Thap11*<sup>dTAG/dTAG</sup> (dTAGv1-Homo) E5.5 epiblast cells. The x and y axis of the dot plots are Log (CPM+1). **b**, Heatmap showing the expression of Epi-, ExE-, and VE-marker genes across single-cell RNA-seq datasets of Epi, ExE,

and VE <sup>2</sup>, together with our bulk low-input total RNA-seq data from Epi samples. **c**, Heatmap showing the expression of naïve- and formative-pluripotency-marker genes based on bulk low-input total RNA-seq data of Epi samples. **d**, Schematic diagram of experimental design. **e**, Heatmap showing the expression dynamics of naïve- and formative-pluripotency-marker genes in 3D-cultured THAP11-dTAG ESCs with DMSO or dTAG13 treatment during induction. **f**, Schematic diagram of experimental design. **g-h**, Expression dynamics of naïve- and formative-pluripotency-marker genes in 2D-cultured THAP11-dTAG ESCs with DMSO or dTAG13 treatment during EpiLC differentiation. **i**, Scatter plots comparing gene expression profiles of E5.5 Epi cells from DMSO-Homo, dTAGv1-Het, and dTAGv1-Homo embryos. The x and y axis of the dot plots are Log2CPM (counts per million) from RNA-seq. Fold change > 2, false discovery rate (FDR) < 0.05.

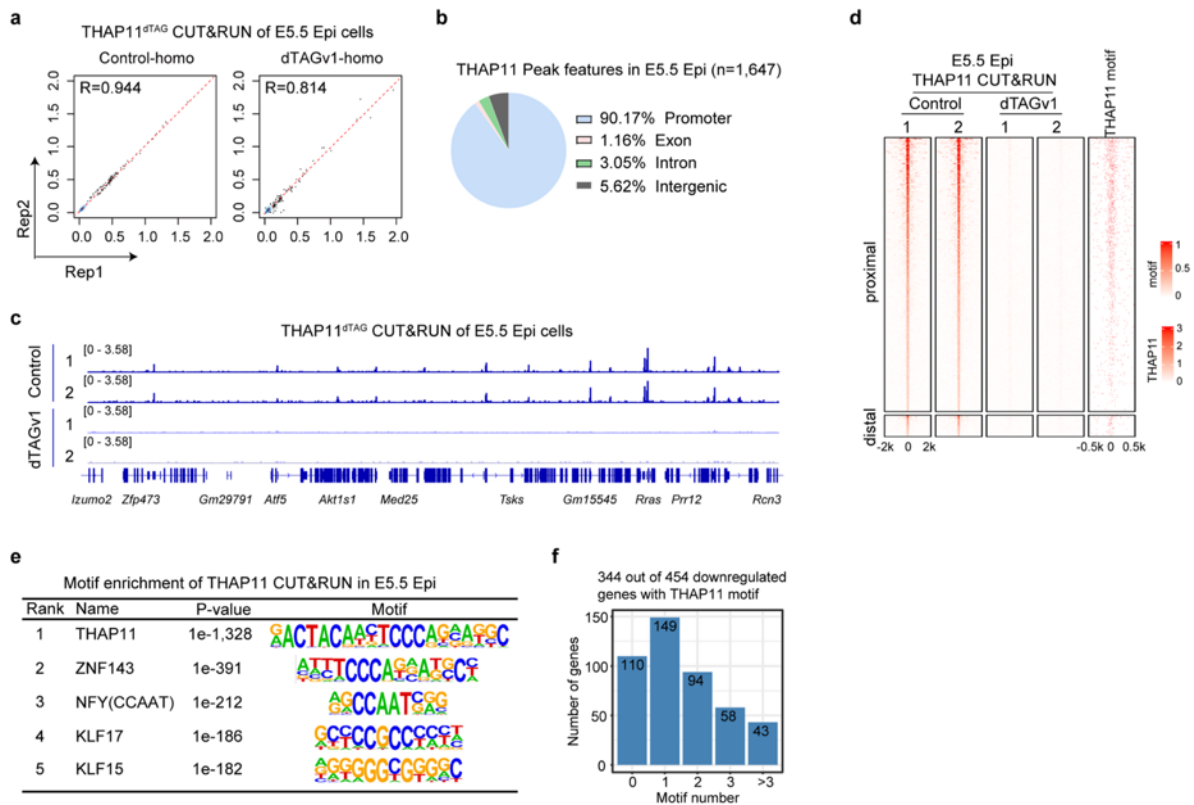

**Supplementary information, Fig. S3. THAP11 regulates polarity genes through promoter binding in epiblast cells.** **a**, Correlation of replicates for THAP11 CUT&RUN in E5.5 Epi cells. Each dot represents a 5-kb bin. **b**, The genomic distribution of THAP11 binding peaks generated by THAP11 CUT&RUN in E5.5 Epi cells. **c**, Genome browser view showing examples of THAP11 CUT&RUN profiles in E5.5 Epi cells with or without dTAGv1 treatment. **d**, Heatmaps showing all the CUT&RUN signals of THAP11 peaks at promoters and distal regions in E5.5 Epi cells with or without dTAGv1 treatment. **e**, The top 5 TF motifs enriched in the regions bound by THAP11 in E5.5 Epi cells. Note that based on the p-value, the THAP11 motif is much more highly enriched compared to the other enriched motifs. **f**, THAP11 motif numbers found in the promoters of the 454 downregulated genes (dTAGv1-Homo VS DMSO).

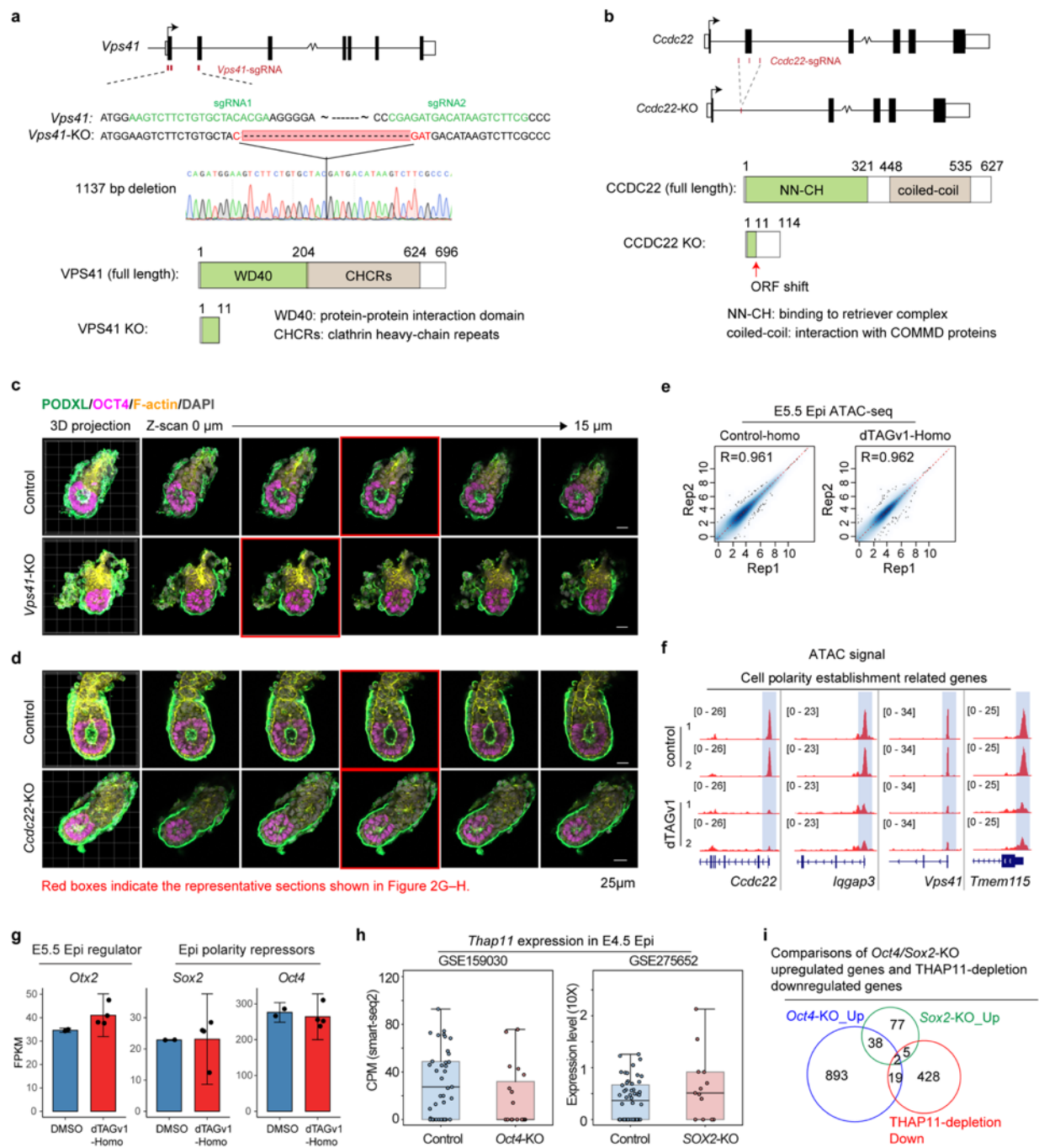

**Supplementary information, Fig. S4. *Vps41* and *Ccdc22* knockout impairs Epi lumenogenesis.**

**a**, Schematic diagrams illustrating the *Vps41* knockout strategy, in which CRISPR-mediated deletion of exons 1–2, encoding the WD40 domain, results in premature translation termination.

**b**, Schematic diagrams illustrating the *Ccdc22* knockout strategy, in which CRISPR-mediated deletion of exon 2, encoding the NH-CH domain, results in premature translation termination.

**c–d**, Immunostaining of OCT4, F-actin, and PODXL in control, *Vps41*-KO (c), and *Ccdc22*-KO (d)

E5.5 embryos. The red boxes indicate the representative sections shown in Fig. 2g-h. Scale bar, 25 μm. **e**, Correlation of replicates for ATAC-seq in control or THAP11-depleted E5.5 Epi. The x and y axis of the dot plots are $\log_2(\text{normalized\_counts}+1)$. Each dot represents a 5-kb bin. **f**, Examples of genome browser view of ATAC signals in control and THAP11-depleted E5.5 Epi. **g**, Expression levels of *Otx2*, *Sox2*, and *Oct4* in control or THAP11-depleted E5.5 Epi cells. **h**, Expression levels of *Thap11* in E4.5 *Oct4*-KO (GSE159030) or *Sox2*-KO (GSE275652) Epi. **i**, Venn diagram showing the overlapped genes downregulated in the E5.5 THAP11-depleted Epi cells and upregulated in the E4.5 *Oct4*- or *Sox2*-KO Epi cells.

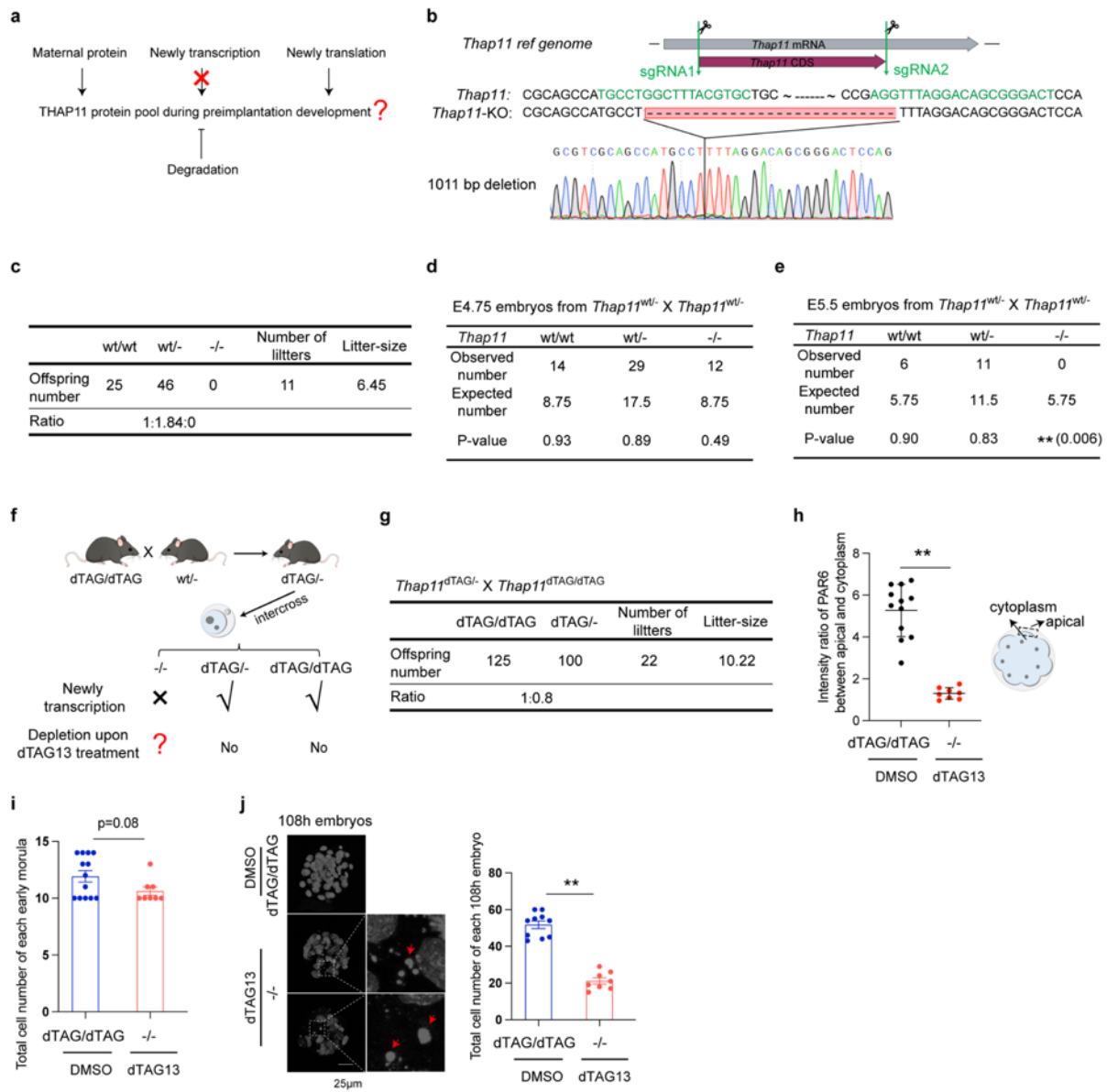

**Supplementary information, Fig. S5. Generation of the THAP11<sup>dTAG/-</sup> mouse line. a,** Schematic diagrams illustrating the THAP11 protein depletion strategy. **b,** Schematic diagrams illustrating the *Thap11* knockout strategy, in which CRISPR-mediated deletion of entire protein coding region. **c,** Statistics of pup numbers from THAP11<sup>wt/-</sup> × THAP11<sup>wt/-</sup> crosses, indicating successful generation of THAP11 knockout allele. **d,** Genotype ratios of E4.75 embryos from *Thap11*<sup>wt/-</sup> intercrosses. Chi-square test. **e,** Genotype ratios of E5.5 embryos from *Thap11*<sup>wt/-</sup> intercrosses. Chi-square test. **f,** The experimental design for the generation of THAP11-depleted embryos. **g,** Statistics of pup numbers from THAP11<sup>dTAG/-</sup> × THAP11<sup>dTAG/dTAG</sup> crosses, indicating normal function of the THAP11<sup>dTAG</sup> fusion protein. **h,** Intensity ratio of PAR6 between apical and

cytoplasm in control (n=12) or THAP11-depleted (n=8) early morula. $^{**}p < 0.01$, Student's t test.
**i**, Total cell number of control (n=13) and THAP11-depleted (n=8) early morulae. **j**, left panel:
representative image of control and THAP11-depleted embryos at 108h. Scale bar, 25 μm. Red
arrowheads indicate DNA fragmentation. Right panel: Total cell number of control (n=10) and
THAP11-depleted (n=8) embryos at 108h. $^{**}p < 0.01$, Student's t test.

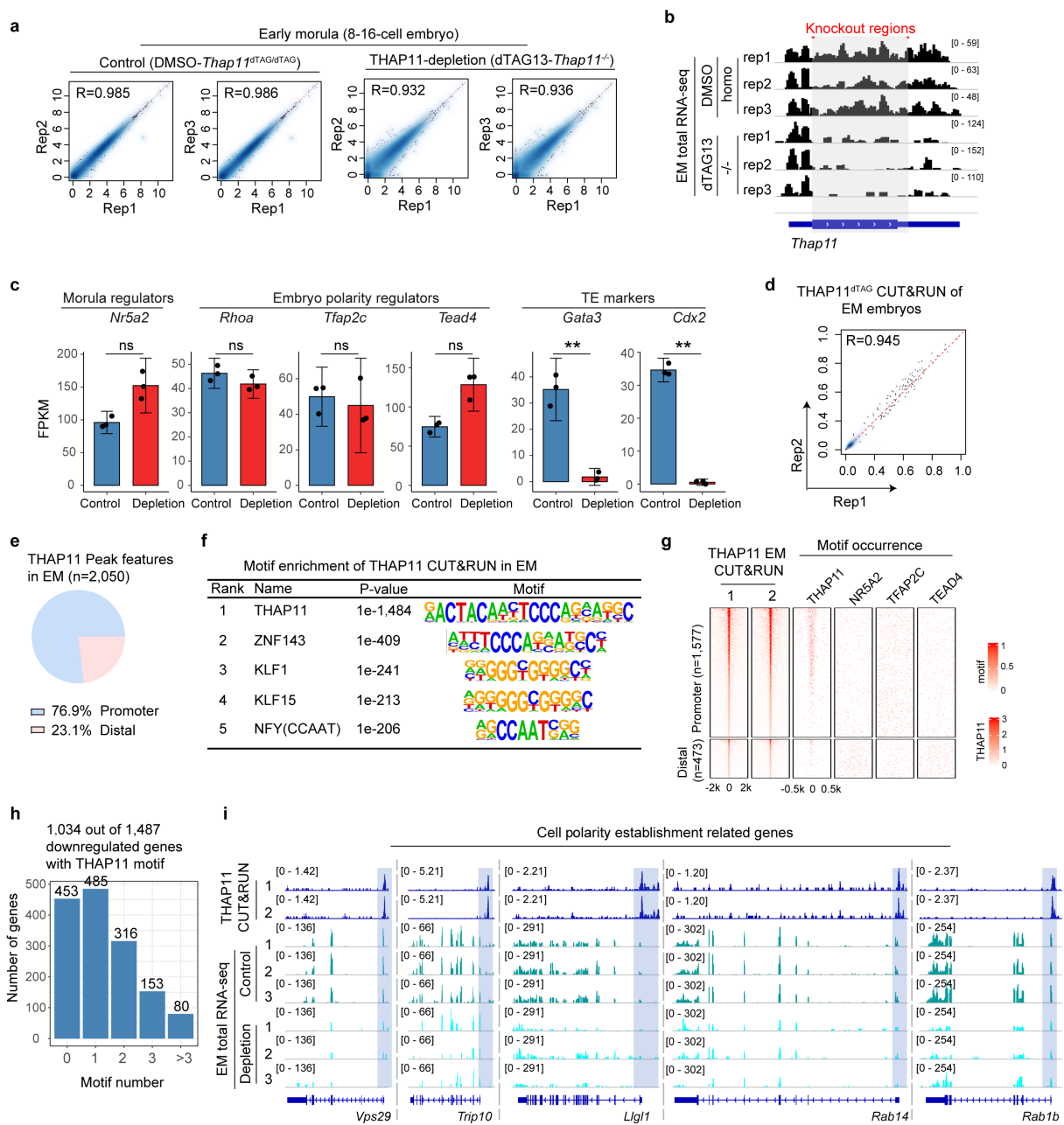

**Supplementary information, Fig. S6. THAP11 regulates polarity genes through promoter binding in EM.** **a**, Correlation of replicates for total RNA-seq in EM of control (DMSO-*Thap11*<sup>dTAG/dTAG</sup>) and THAP11-depletion (dTAG13-*Thap11*<sup>-/-</sup>). The x and y axis of the dot plots are Log (CPM+1). **b**, Genome browser view of RNA-seq at the *Thap11* genomic regions in EM of control and THAP11-depletion. **c**, Expression levels of *Nr5a2*, *Rhoa*, *Tfap2c*, *Tead4*, *Gata3*, and *Cdx2* in control or THAP11-depleted EM. \*\*, FDR<0.01; ns, FDR>0.05. **d**, Correlation of replicates for THAP11 CUT&RUN in EM. Each dot represents a 5-kb bin. **e**, The genomic

distribution of THAP11 binding peaks generated by THAP11 CUT&RUN in EM. **f**, The top 5 TF
motifs enriched in the regions bound by THAP11 in EM. Note that based on the p-value, the
THAP11 motif is much more highly enriched compared to the other enriched motifs. **g**, Heatmaps
showing all the CUT&RUN signals and motif occurrence of the THAP11 peaks at promoters and
distal regions in EM. **h**, THAP11 motif numbers found in the promoters of the 1,487
downregulated genes without detectable CUT&RUN signal upon THAP11-depletion. **i**, Genome
browser view of THAP11 targeted polarity gene examples showing THAP11 CUT&RUN and
RNA-seq in EM.

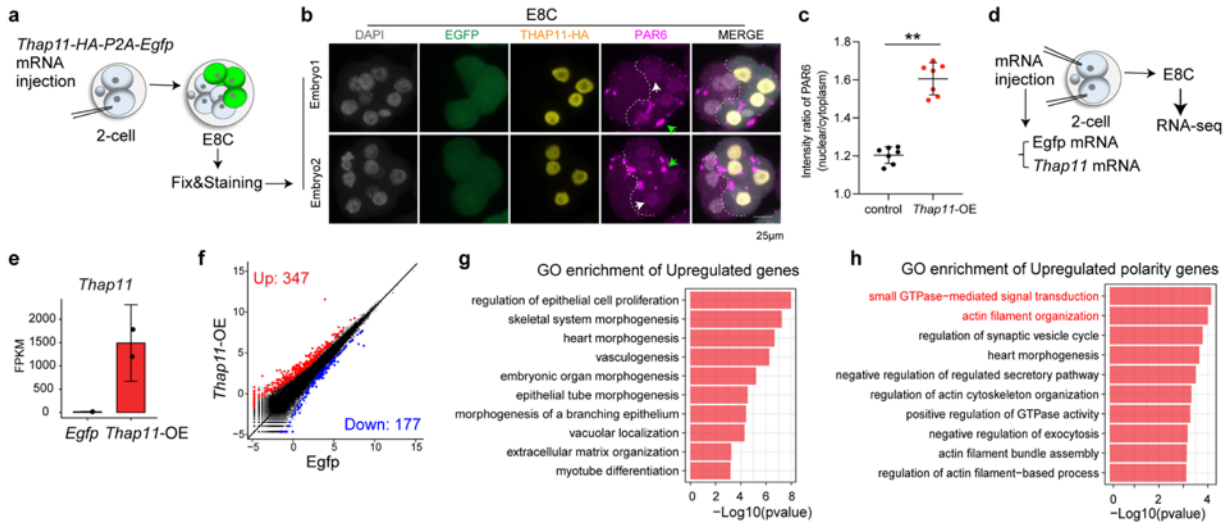

**Supplementary information, Fig. S7. THAP11 overexpression promotes premature clustering of apical proteins and activation of important polarity program.** **a**, Schematic diagram of experimental design. One blastomere of late 2-cell stage embryos was injected with Thap11-HA-P2A-Egfp mRNA, and the embryos were subsequently collected at early 8-cell (E8C) stage for immunostaining. **b**, Immunostaining of HA (THAP11-HA) and PAR6 in E8C embryos. White arrows indicate nuclear PAR6. Green arrows indicate membrane large PAR6 cluster. **c**, Intensity ratio of PAR6 between nuclear and cytoplasm in control or THAP11-OE blastomeres. Data are presented as mean ± SEM; \*\*$p < 0.01$, Student's t test. **d**, Schematic diagram of experimental design. Late 2-cell stage embryos were injected with either Egfp- or Thap11-HA-P2A-Egfp mRNA, and embryos were subsequently collected at E8C stage for RNA-seq. **e**, Expression of Thap11 in Egfp- or Thap11-OE E8C embryos. **f**, Scatter plots comparing the gene expression profiles of E8C embryos with *Egfp* or *Thap11* overexpression. The x and y axis of the dot plots are Log<sub>2</sub>CPM from RNA-seq. Fold change $> 2$, FDR $< 0.05$. **g**, Representative GO terms enriched in the upregulated genes (n=347) in response to *Thap11* overexpression. **h**, Representative GO terms enriched in the upregulated polarity genes in response to *Thap11* overexpression.

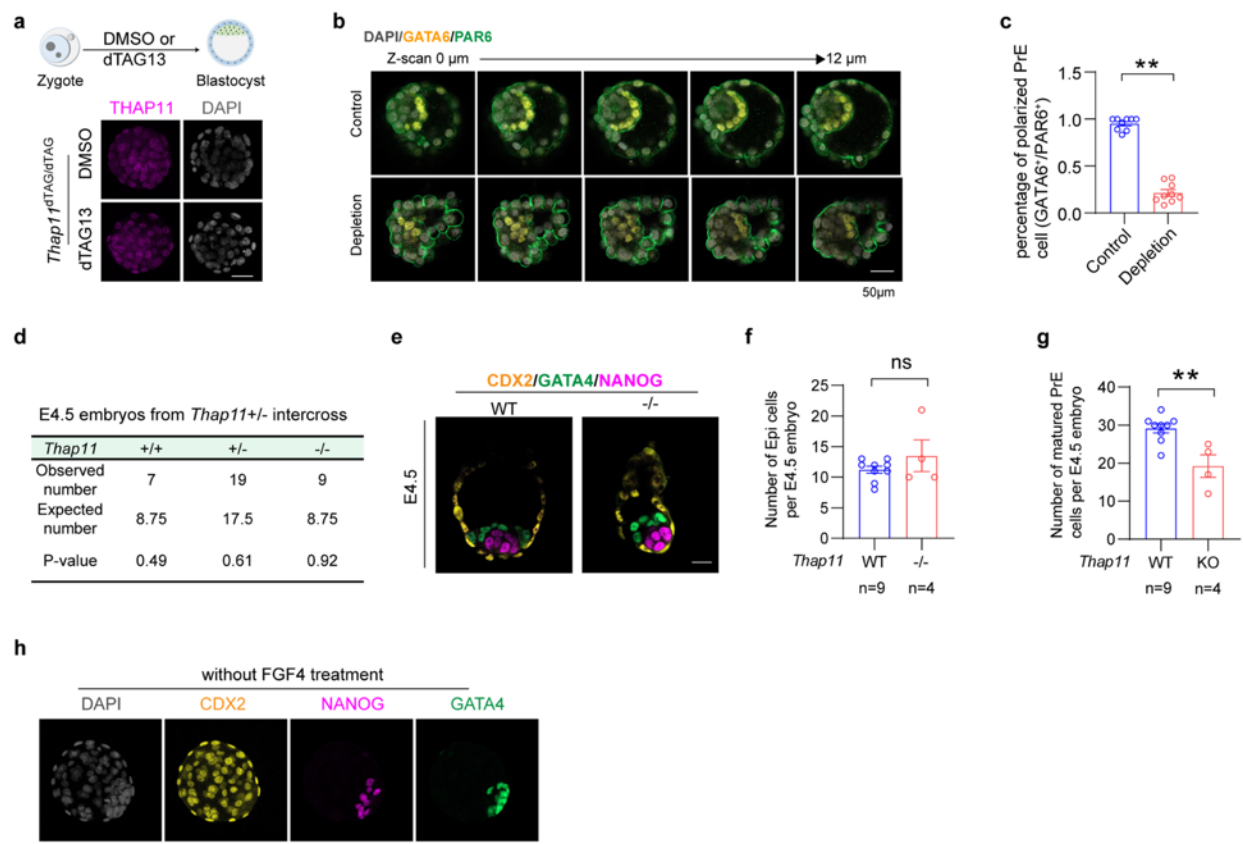

**Supplementary information, Fig. S8. THAP11 depletion causes failure of PrE polarization and maturation.** **a**, Immunostaining of THAP11 in *Thap11*<sup>dTAG/dTAG</sup> blastocysts with or without dTAG13 treatment. **b**, Individual Z-stack optical sections of GATA6 and PAR6 immunostaining in control and THAP11-depleted LB. The corresponding maximum-intensity projection is shown in Fig. 5b. **c**, Percentage of polarized PrE cells (GATA6<sup>+</sup>/PAR6<sup>+</sup>) per embryos of control and THAP11-depletion. Mean ± SEM; p-value, Student's t test. \*\*, p<0.01. **d**, Genotype ratios of E4.5 embryos from *Thap11*<sup>+/-</sup> intercrosses. Chi-square test. **e**, Immunostaining of lineage markers CDX2, GATA4, and NANOG in WT and *Thap11*<sup>-/-</sup> E4.5 embryos. **f**, Number of Epi cells per E4.5 embryos of WT and *Thap11*<sup>-/-</sup>. Mean ± SEM; p-value, Student's t test. ns, p>0.05. **g**, Number of matured PrE cells per E4.5 embryos of WT and *Thap11*<sup>-/-</sup>. **h**, Immunostaining of lineage markers CDX2, NANOG, and GATA4 in WT LB embryos without FGF4 treatment. Mean ± SEM; p-value, Student's t test. \*\*, p<0.01. Scale bar, 50 μm.

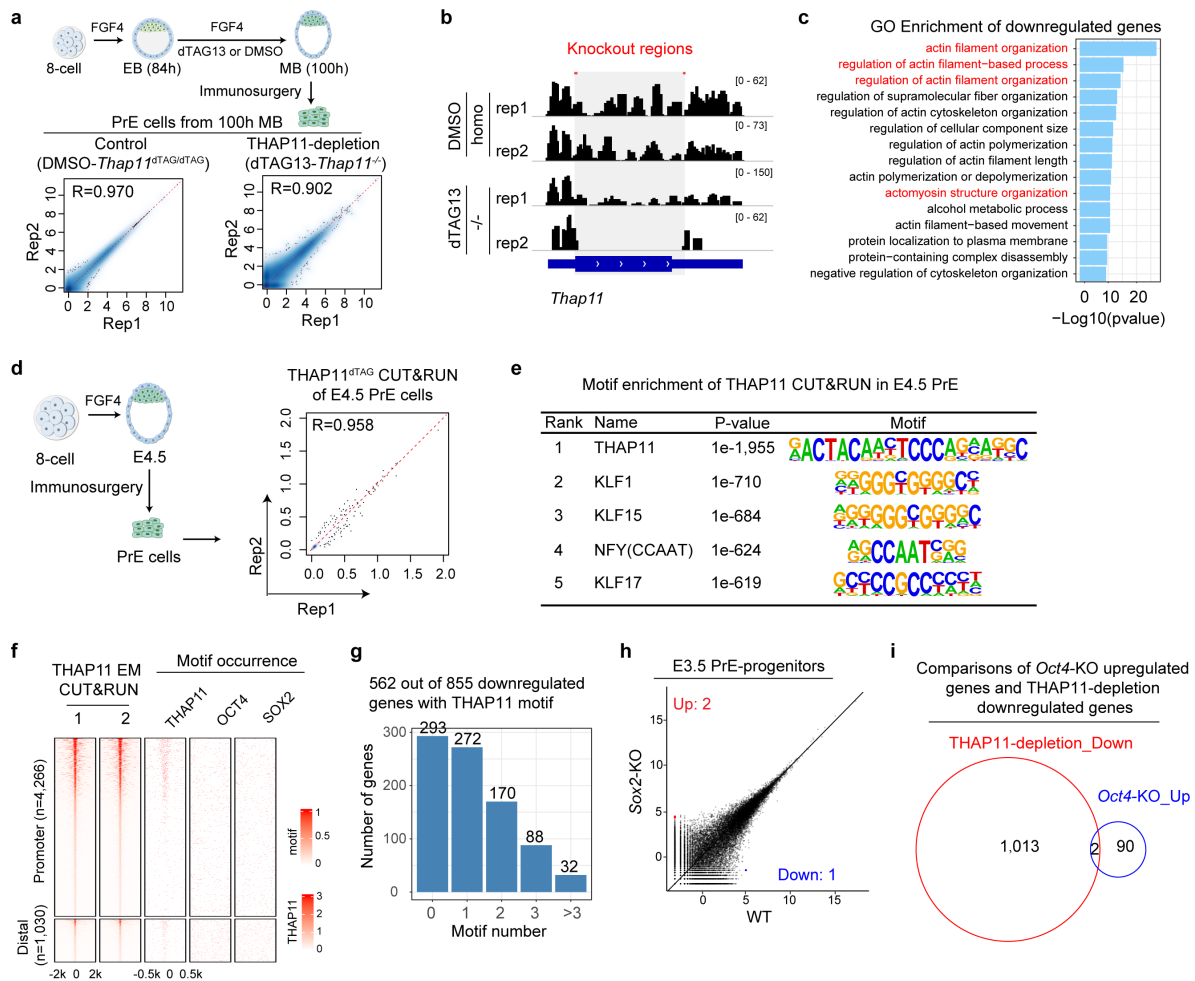

**Supplementary information, Fig. S9. THAP11 regulates polarity genes through promoter binding in PrE cells.** **a**, Upper panel: schematic diagram of experimental design. Lower panel: correlation of replicates for total RNA-seq in PrE cells of control (DMSO-*Thap11*<sup>dTAG/dTAG</sup>) and THAP11-depletion (*dTAG13-Thap11*<sup>-/-</sup>). The x and y axis of the dot plots are Log (CPM+1). **b**, Genome browser view of RNA-seq at the *Thap11* genomic regions in PrE cells of control and THAP11-depletion. **c**, GO enrichment of downregulated genes (n=1,015) in response to THAP11-depletion in PrE cells. **d**, Left panel: schematic diagram of experimental design. Right panel: Correlation of replicates for THAP11 CUT&RUN in E4.5 PrE cells. Each dot represents a 5-kb bin. **e**, The top 5 TF motifs enriched in the regions bound by THAP11 in PrE cells. Note that based on the p-value, the THAP11 motif is much more highly enriched compared to the other enriched motifs. **f**, Heatmaps showing all the CUT&RUN signals and motif occurrence of THAP11 peaks at promoters and distal regions in PrE cells. **g**, THAP11 motif numbers found in the promoters of the other 855 downregulated genes without detectable CUT&RUN signal upon THAP11-depletion.

**h**, Scatter plots comparing the gene expression profiles of E3.5 PrE progenitors upon *Sox2*-KO.
162 The x and y axis of the dot plots are Log<sub>2</sub>CPM from RNA-seq. Fold change > 2, FDR < 0.05. **i**,
163 Venn diagram showing the overlapped genes between downregulated in THAP11-depleted PrE
cells and upregulated in *Oct4*-KO PrE progenitors.

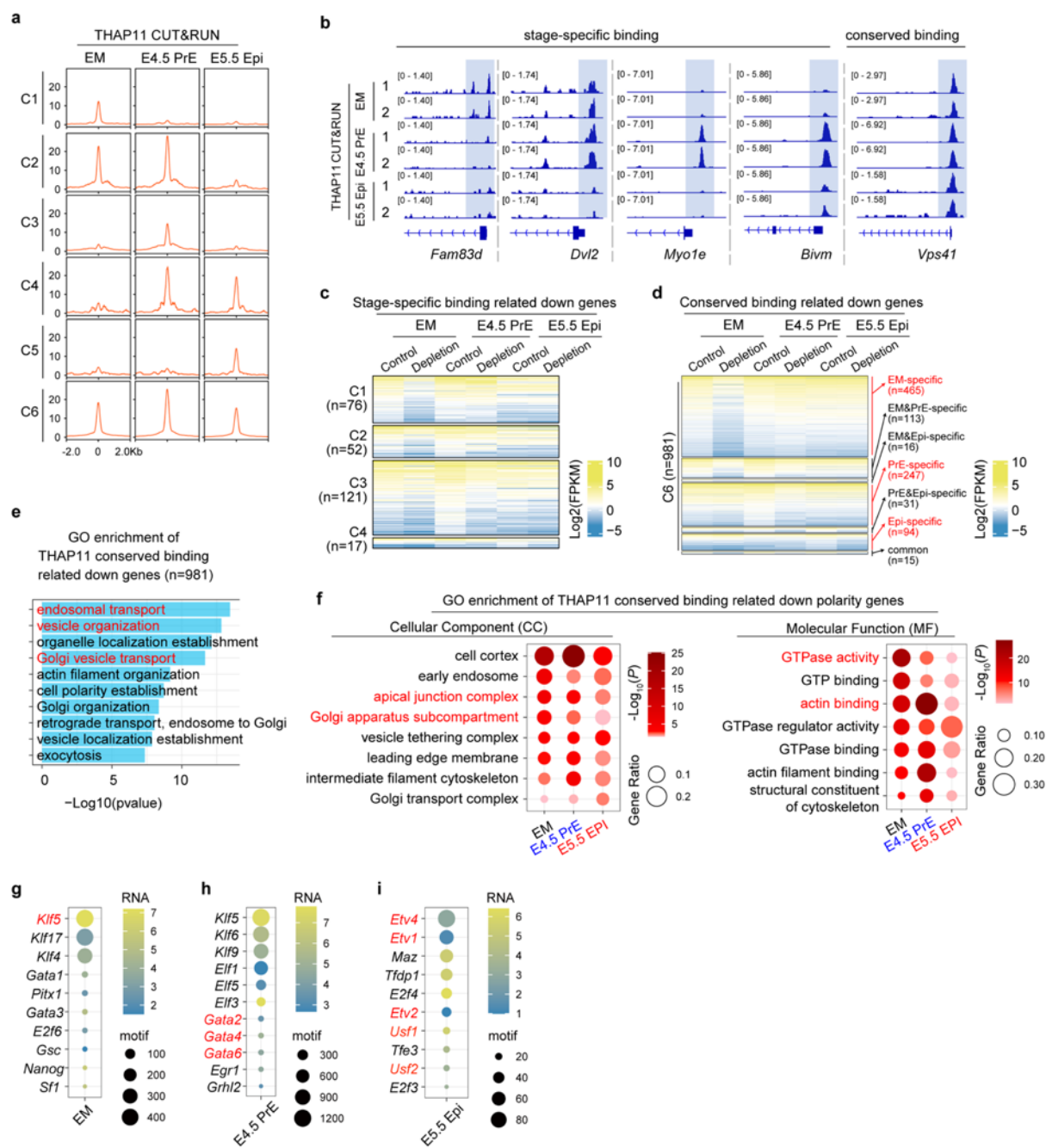

**Supplementary information, Fig. S10. Common and stage-specific THAP11 chromatin bindings regulate polarity programs.** **a**, Density plots of different THAP11 binding clusters (C1-C6). **b**, Genome browser view of THAP11 stage-specific and conserved targets showing CUT&RUN at the indicated stages. **c**, Heatmaps showing the gene expression of the stage-specific binding (C1-C5) associated downregulated genes (FDR<0.05, LogFC<0) in EM, E4.5 PrE, and E5.5 Epi upon THAP11 loss. **d**, Heatmaps showing the gene expression of the common binding

(C6) associated downregulated genes (FDR<0.05, LogFC<0, n=981) in EM, E4.5 PrE, and E5.5
Epi upon THAP11 loss. These genes were further sub-grouped into EM-specific (n=465),
EM&PrE-specific (n=113), EM&Epi-specific (n=16), PrE-specific (n=247), PrE&Epi-specific
(n=31), Epi-specific (n=94), and common (n=15) groups. **e**, Representative GO terms enriched
among THAP11 conserved binding-related downregulated genes (n=981). **f**, Representative GO
(CC and MF) terms enriched among THAP11 conserved binding-related downregulated polarity
genes at indicated stages. **g-i**, Enrichment and expression levels of top TF motifs at the THAP11
CUT&RUN peaks of EM (g), E4.5 PrE (h), and E5.5 Epi (i).

**Supplemental tables (separated files)**

**Supplementary Table 1:** Polarity gene lists used in this study

**Supplementary Table 2:** List of differentially expressed genes in E5.5 Epi in response to THAP11 depletion

**Supplementary Table 3:** List of differentially expressed polarity genes in Epi of THAP11-depletion and *Oct4*- or *Sox2*-KO

**Supplementary Table 4:** List of differentially expressed genes in EM of THAP11-depletion versus control

**Supplementary Table 5:** List of differentially expressed polarity genes in EM of THAP11-depletion and *Tfap2c*/*Tead4*-KD

**Supplementary Table 6:** List of differentially expressed polarity genes in E8C of THAP11-overexpression

**Supplementary Table 7:** List of differentially expressed genes in PrE of THAP11-depletion versus control

**Supplementary Table 8:** List of differentially expressed genes in E3.5 PrE progenitors of *Oct4*- or *Sox2*-KO versus control

**Supplementary Table 9:** List of differentially expressed polarity genes in PrE of THAP11-depletion and *Oct4*- or *Sox2*-KO.

**Supplementary Table 10:** List of THAP11 stage-specific binding related downregulated genes.

**Supplementary Table 11:** List of THAP11 common binding related downregulated genes at indicated stages.

**Supplementary Table 12:** Candidate cofactors that may cooperate with THAP11

**Supplementary Table 13:** List of primers used in this study

**Supplementary Table 14:** ddPCR validated THAP11-dTAG knock-in copy number

**Supplementary Table 15:** Summary of the sequenced libraries in this study

**Supplementary Table 16:** Summary of the public datasets used in this study
